# Structures of Cas9-Bound Double-Stranded DNA Mini-Circle Reveal Impacts of DNA Shape on Cas9 Target Interrogation

**DOI:** 10.64898/2026.05.20.726649

**Authors:** Kyu-Yeon Lee, Donggyun Kim, Yukang Liu, Hongjian Chang, Vadim Cherezov, Jiansen Jiang, Peter Z. Qin

## Abstract

CRISPR-Cas9 is an RNA-guided endonuclease that cleaves double-stranded DNA at specific sites and has been adapted as a powerful tool for genome manipulation. Cas9 recognizes its target through multiple conformational transitions coordinated between the Cas9 ribonucleoprotein and the DNA duplex. Such transitions, and consequently Cas9 targeting specificity, are expected to be significantly influenced by the collective duplex physical properties referred to as DNA shape. To advance our currently limited understanding of the interplay between DNA shape and Cas9 target interrogation, we solved two cryo-EM structures of SpyCas9 bound to a cognate target embedded in a relaxed 95-base-pair DNA double-stranded minicircle. The Cas9-bound DNA segment engages in similar interactions involved in PAM-binding and R-loop initiation as those observed in Cas9-bound linear DNA. However, R-loop is limited to less than three base-pairs, thus interfering with Cas9 cleavage. The minicircle DNA, which is fully resolved, retains its global shape. As Cas9 locally unwinds the protospacer, the closed-ring topology constrains the movement of the paired PAM-distal DNA duplex, thus interfering with R-loop propagation. These data provide detailed insight into the interplay between DNA shape and Cas9 structure and function, and may shed light on genome-editing and manipulation in environments with varied DNA topologies.

## Introduction

CRISPR-Cas (clustered regularly interspaced short palindromic repeats and CRISPR-associated proteins) nucleases natively function as an integrated component of RNA-guided adaptive immune systems in bacteria and archaea to provide protection against invasion of foreign genetic elements.^1^ Understanding the mechanism of CRISPR function has enabled its broad adaptation for genome manipulation and engineering, unleashing a technological revolution that continues to advance rapidly.^2–4^ Among the large number of CRISPR nucleases discovered, Cas9, a member of the Class 2, type II CRISPR family,^1,5^ has become the workhorse for genome manipulation.^2–4^ Cas9 cleaves double-stranded DNAs in a sequence-specific manner using a protein/RNA effector complex (i.e., the enzyme) where the RNA can be engineered as a single-guide-RNA (sgRNA).^6–9^ The Cas9 effector enzyme orchestrates a series of complex and coordinated conformational changes to identify its cognate DNA target.^10–13^ More than a decade of extensive studies have identified the major check-points within the general mechanistic framework of Cas9 target interrogation, which include:^10–13^ (1) initial DNA recognition by effector binding to the protospacer adjacent motif (PAM), one of the two essential elements of a target; (2) bending of PAM-adjacent duplex to initiate R-loop formation, where the single-stranded guide of sgRNA base pairs with the target-strand (TS) of the DNA protospacer; (3) further protospacer unwinding and propagation of the R-loop toward the PAM-distal segment, which relies on complementarity of the RNA guide and TS; and (4) additional R-loop dependent protein conformational changes to enable HNH nuclease domain cleavage of TS and RuvC nuclease domain cleavage of the other strand (designated as the non-target-strand, NTS).

Advances in mechanistic understanding of Cas9 target discrimination have led to technological breakthroughs.^4^ For example, a number of Cas9 variants have been engineered to achieve enhanced specificity toward off-target sites containing mismatches between the DNA protospacer and the guide RNA.^14–19^ However, our current mechanistic understanding is not yet complete. One of the major remaining knowledge gaps is how the collective physical properties of a DNA duplex, referred to as DNA shape, impact Cas9 target interrogation. DNA shape encompasses a wide-range of physical properties (e.g. flexibility, groove width, local electrostatics) and is determined collectively by the local “core” base-pair(s) and other factors such as (distal) peripheral sequences and topological constraints (e.g., supercoiling).^20,21^ It is well established that DNA shape plays a key role in specific recognition by proteins (e.g., transcription factors) and small-molecules.^20,21^ Because a DNA duplex undergoes extensive distortions upon interacting with Cas9,^10–13^ DNA shape is expected to be a significant factor in Cas9 target interrogation.

The interplay between DNA shape and Cas9 target interrogation has been explored in a number of studies. For example, DNA supercoiling has been reported to influence PAM recognition,^22,23^ accelerate Cas9 nicking^24,25^ and double-strand-break with cognate targets,^24–28^ alter off-targets cleavage,^29–31^ and promote DNA turnover.^28^ In addition, with truncated guides that are shorter than the normal full-length of 20-nucleotide (-nt), DNA strand scission is modulated by both negative supercoiling as well as the intrinsic DNA duplex dissociation energy at the protospacer segment beyond the RNA/DNA hybrid.^25,32^ However, as the vast majority of Cas9 studies use linear DNA substrates, investigations into the impact of DNA shape remain in the early stages.

Minicircles are circular double-stranded DNA that form a closed loop with no ends.^33^ Beyond the primary sequence, the additional topological constraints of minicircles provide an attractive platform for studying DNA shape and its impact on recognition by proteins and other partners.^31,34,35^ Recently, we have constructed a 95 base-pair (bp) double-stranded DNA minicircle (dsMC95). A 5.3 Å cryo-EM structure of the free dsMC95 shows that the duplexed DNA forms a relaxed circle with modest ellipticity, minor out-of-plane displacement, and no observable local deformation.^36^ The in-plane dimension of dsMC95,which correlates with bending of the duplex, is comparable to DNA present in core nucleosome particles.^36^ Although the duplex retains a B-DNA conformation, it differs from linear DNA by exhibiting widened outward-facing major grooves and compressed inward-facing ones.^36^

Building on the understanding of the intrinsic DNA shape of dsMC95, here we present cryo-EM studies of dsMC95 bound by a *Streptococcus pyogenes* Cas9 ribonucleoprotein (RNP) complex (designated as “Cas9” from here on). We were able to reconstitute two atomic resolution maps at resolutions of 2.7 Å and 3.1 Å, respectively, from a single cryo-EM dataset. The Cas9-dsMC95 complexes maintain similar interactions involved in PAM-binding and R-loop initiation as those observed in structures of Cas9-bound linear DNA. However, unlike the recently reported structures of Cas9 bound to negatively supercoiled 126-bp minicircles that show complete R-loop formation,^31^ the relaxed dsMC95 minicircle stalls R-loop propagation and enables capturing of early R-loop intermediates, including a 3-bp R-loop and previously unobserved states featuring a nascent 2-bp and a 1-bp R-loop. Our analysis shows that Cas9 acts as a rigid-body to search for PAM and initiate R-loop formation with the relaxed dsMC95 minicircle, which is the same mechanism used for linear or negatively supercoiled substrates. Comparisons of the completely resolved dsMC95 minicircles show that the DNA maintains its global shape but deforms locally at the Cas9 interacting segment. The close-ring topology of the relaxed dsMC95 minicircle constrains the PAM-distal paired DNA movements towards the pre-positioned RNA guide, thus interfering with R-loop propagation and DNA cleavage. The results unambiguously demonstrate that Cas9 function can be affected by intrinsic DNA shape features, in this case the relaxed circle topology, resulted from sequences extending well beyond the PAM and the 20-bp protospacer. These insights not only advance our fundamental understanding of the CRISPR-Cas9 mechanism but also have practical implications for predicting and optimizing genome-editing efficiency in complex, topologically constrained genomic environments.

## Results and Discussion

### A dCas9-bound relaxed 95-basepair minicircle adopted a multitude of states

To investigate how DNA topology influences R-loop formation and Cas9 recognition of DNA targets, we carried out cryo-EM studies of a relaxed 95-bp double-stranded DNA minicircle (dsMC95) bound by a catalytically-inactive Cas9 (dCas9) (Figure 1). dsMC95 was synthesized using a dumbbell-mediated ligation method,^36^ and contained a canonical Cas9 target with a 20-nt protospacer upstream of a TGG PAM [Supplementary Information (SI), Figure S1, Table S1]. dsMC95 was subjected to binding by a dCas9 RNP containing a single-guide RNA (sgRNA) with a 20-nt guide that fully matches the protospacer (Figure 1a, 1b; Table S1). The ternary dCas9-sgRNA-dsMC95 complex was isolated via size exclusion chromatography (Figure S2). Biochemical characterizations showed that Cas9 can bind to dsMC95 (Figure S3) and induce DNA unwinding within the protospacer (Figure S4). However, Cas9 binding to dsMC95 was significantly weaker compared to that of a linear DNA duplex (Figure S3), and no dsMC95 cleavage was observed (Figure S5).

**Figure 1.**
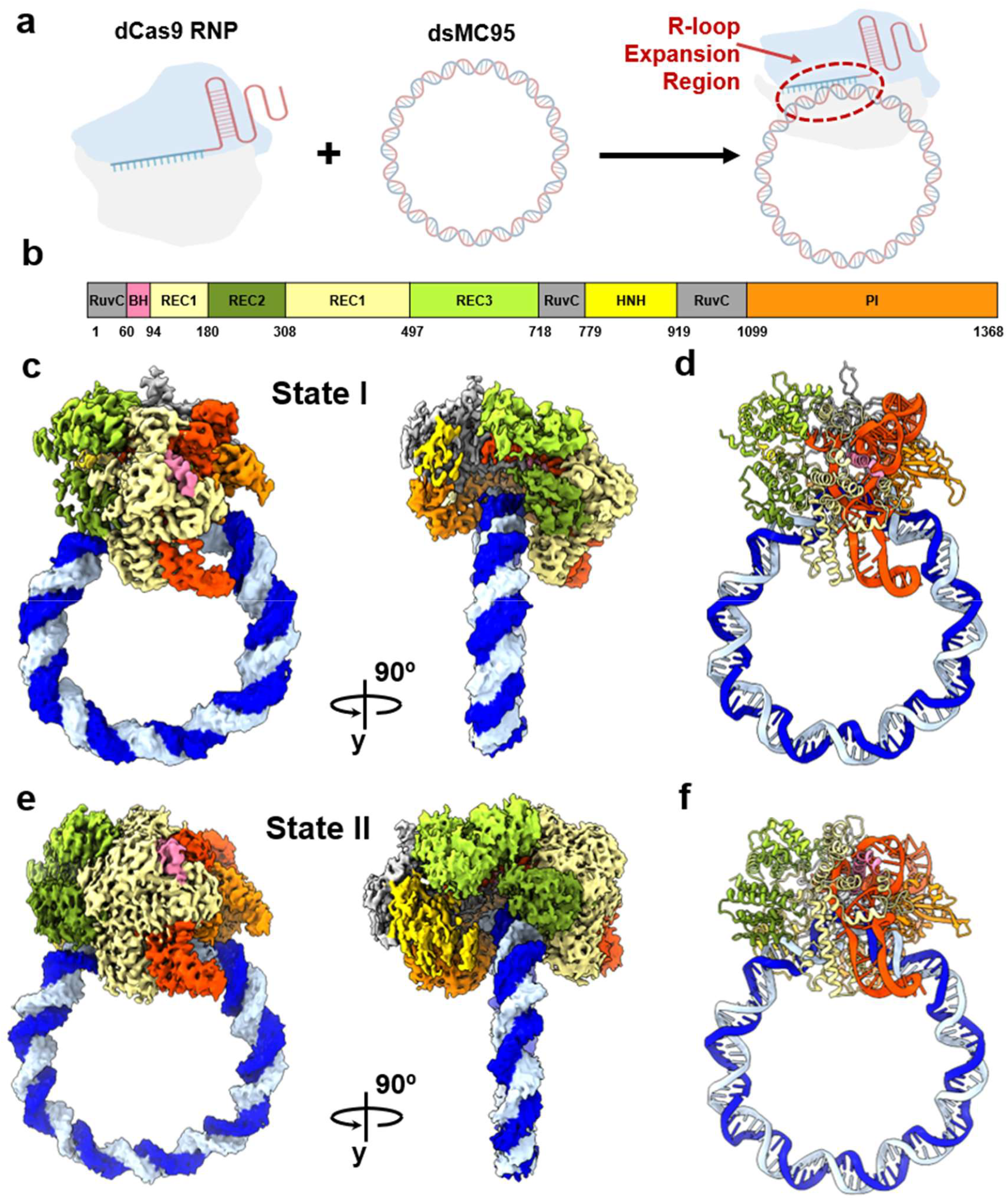
Cryo-EM structures of Cas9-bound relaxed 95-basepair DNA minicircles. **a**, Schematic of assembly of the ternary dCas9-sgRNA-dsMC95 complex. The red dashed oval indicates the PAM-extension region. **b**, Domain organization of *S. pyogenes* Cas9. Domains are colored as follows: RuvC (grey), BH (pink), REC1–3 (green to light yellow), HNH (yellow), and PI (orange). Numbers indicate residue numbers. **c**, Cryo-EM density map of State I. Front and side views are shown with sharpened map at a contour level of 1.48. **d**, Fitted atomic model of State I. **e**, Cryo-EM density map of State II. Front and side views are shown with sharpened map at a contour level of 1.61. **f**, Fitted atomic model of State II. In panels **c**-**f**, DNA strands are shown in dark blue (TS) and light blue (NTS), sgRNA is shown in orange, and the Cas9 protein domains are colored coded as those in panel **b**. See additional information in SI.

Cryo-EM data were acquired on the ternary complex sample using a 300 kV transmission electron microscope Krios equipped with a Gatan BioQuantum K3 Energy Filter and a Gatan K3 direct-electron detector (see Methods). Consistent with the observed disruption of Cas9 binding and cleavage (Figures S3, S5), the cryo-EM data revealed a high degree of heterogeneity present in the ternary complex sample (Figure S6). The work reported here focuses on the analysis of the 1-to-1 complex of dsMC95 and dCas9 RNP (Figures S7, S8). Other species, such as the unbound DNA and the higher-order complexes (Figure S6), will be subjected to future investigation.

Analysis of particles of the 1-to-1 complex revealed several conformations of the dCas9-bound dsMC95 complex (Figure S7). Two of these conformations, designated as State I and State II, were resolved to a global resolution of 3.1 Å and 2.7 Å, respectively (Figures 1c–1f; Figures S7, S8). As described below, State I captures previously unobserved structures of nascent R-loop intermediates, in which the PAM+1 position forms a fully established RNA/DNA hybrid pair, while the PAM+2 position adopts dual conformations, including both DNA/DNA base pairing and RNA/DNA hybrid pairing (Fig. 1c, 1d; Figure S9). State II represents a more advanced intermediate with a stable 3-bp R-loop (Fig. 1e, 1f; Figure S10).

In addition to State I and State II, other conformational classes of the 1-to-1 complex were observed, but low-resolution maps precluded model building (Figure S7). 2-amino-purine fluorescence measurements indicated that dCas9 binding induced a certain degree of DNA unwinding at the PAM+14 position of the protospacer (Figure S4). It is possible that the unresolved conformations present in the 1-to-1 complex dataset represent complexes with different and perhaps more extended R-loop. This would be of interest in future studies.

### State I captures novel structures of early R-loop intermediates

In State I, dCas9 is well-resolved throughout most of the complex (Figure 2a), with 88% of the protein modeled (1,205 of 1,368 residues). The remaining unresolved segments correspond to the HNH domain (residues 766–837, 848–861, and 872–884) that is known to adapt variable conformations.^10–13^ Furthermore, 90% of the sgRNA (90 of 100 nucleotides) is well-defined (Figure S9). Notably, 12 nucleotides of the RNA guide (nucleotide 9–20) are resolved at a local resolution of 2.5 Å (Figure S9a), enabling detailed structural analysis of State I and its comparison to others. While the overall dsMC95 DNA map resolution is ∼6.6 Å (Figure S8), the region from PAM-7 to +14 (see Figure S1 for nomenclature referencing to PAM), which interacts with the dCas9 RNP, is resolved at a local resolution of 3 Å (Figure S9b). This allowed us to fully model both the target and non-target strands of the entire 95-bp DNA (Figure 1c; Figure S9). The final model of State I, including the circular DNA, achieves a MolProbity score of 1.47 (Table S2).

**Figure 2.**
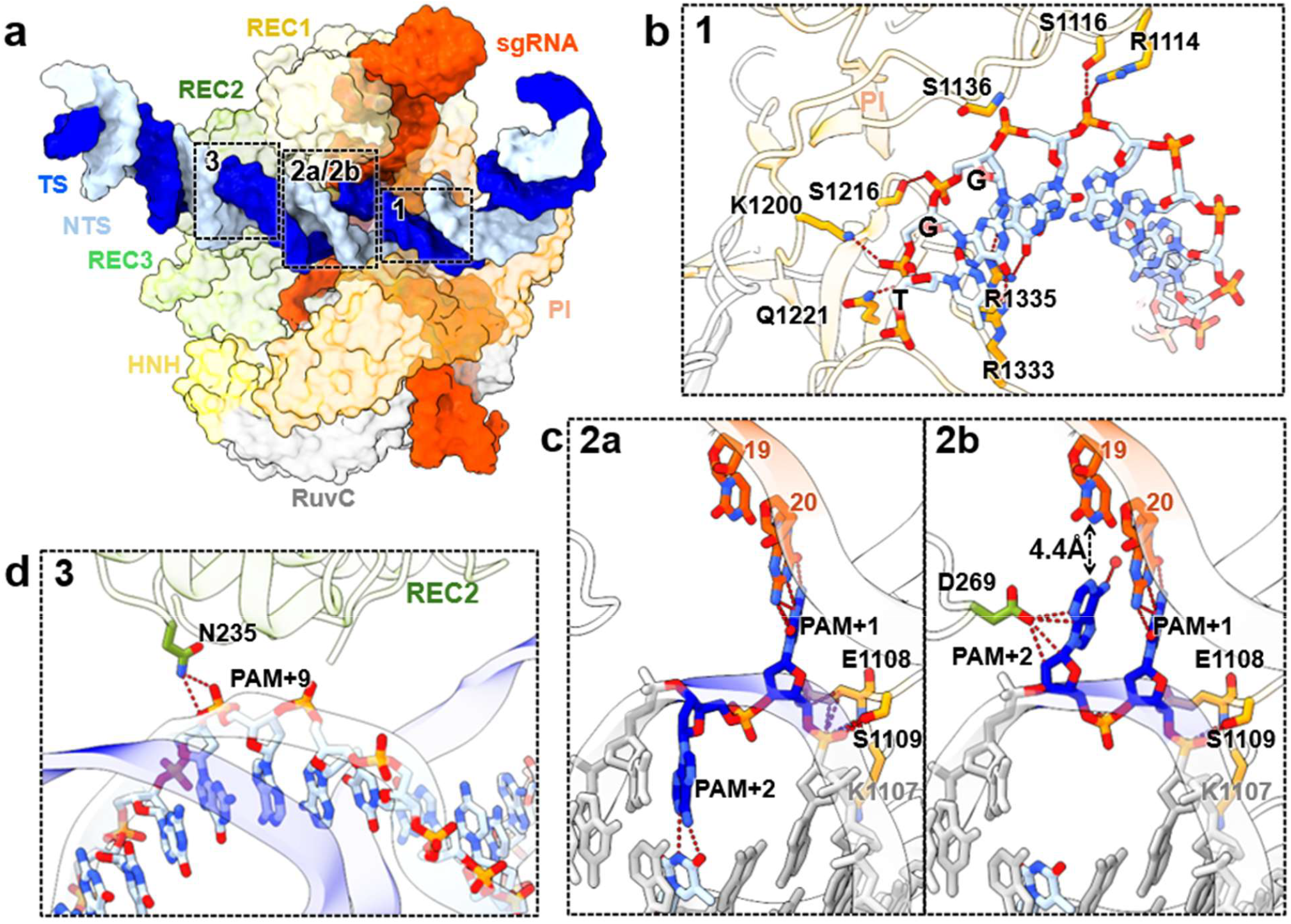
Molecular interactions within State I of the Cas9-dsMC95 complex. **a**, Surface representation of the Cas9-bound DNA segment. Cas9 domains are marked and color-coded as those in Figure 1b. The sgRNA is shown in orange, DNA TS in dark blue and NTS in light blue. Dashed boxes highlight the three key protein–DNA interaction regions with details shown in panels 1–3. **b**, PAM recognition interface. PI-domain residues R1333 and R1335 form base-specific contacts with the PAM sequence (TGG). Additional stabilizing interactions with the DNA phosphate backbone involve residues S1116, R1114, S1136, K1200, S1216 and Q1221. **c**, R-loop initiation at the PAM-proximal region. The phosphate lock (K1107–S1109) interacts with the PAM+1 site. Panel 2a shows the “1-bp R-loop” structure, in which the TS nucleotide at the PAM+2 position remains base-paired with the NTS. Panel 2b shows the “1.5-bp” R-loop structure, where the TS base at PAM+2 breaks DNA-DNA pairing and flips toward the guide RNA nucleotide U19. The flipped PAM+2 TS nucleotide is stabilized by D269 of the REC2 domain and a water molecule. **d**, REC2 interaction with the PAM-distal DNA duplex. Residue N235 from the REC2 domain coordinates the phosphate backbone of the NTS at position PAM+9, likely stabilizing the intact DNA duplex prior to further R-loop propagation. See additional information in SI.

The well-resolved dCas9-bound DNA segment in State I (Figure 2) enables detailed characterization of the similarities and differences between State I and other Cas9 complexes. Structural analysis reveals that the sgRNA conformation and most of the protein-RNA interactions are conserved between State I and the Cas9-sgRNA complex^37^ (PDB id 4ZT0, designated as “binary”), as well as between State 1 and representative ternary complexes assembled with linear DNA substrates (Figure S11). In particular, comparison of State I with the binary complex shows that sgRNA adopts an identical configuration at the PAM-adjacent positions 15-20, with a phosphorus RMSD of 0.67 Å (Figure S11). This indicates that the RNA guide is “pre-folded” into an A-form configuration in the binary complex to “receive” and “interrogate” DNA targets, which is consistent with the established mechanism of Cas9 function.^37^

In State I, the PAM recognition interface closely resembles that of the more advanced R-loop states observed with linear DNA substrates, with all critical interactions between protein side-chains and the DNA PAM segment preserved (Figure 2b, panel 1; Figures S13, S14). This indicates that the topological constraint presented in the relaxed dsMC95 circle does not interfere with PAM engagement. Furthermore, at the PAM+1 position, the phosphate lock is present between K1107 and S1109 and the TS phosphate group, while the TS nucleobase is flipped out of the DNA duplex to form a canonical Watson-Crick (W-C) pairing with the RNA guide (Figure 2c, panel 2a & 2b; Figure S12). This indicates a successful R-loop initiation with the dsMC95 circle.

Within the well-resolved map of State I, the density indicates that the DNA TS nucleotide at the PAM+2 position adopts two distinct conformations (Figure 2c, panel 2a & 2b; Figure S9c). In one of the conformations (Figure 2c, panel 2a), the DNA TS nucleotide (dA66) forms a canonical W-C pair with the NTS PAM+2 nucleotide (dT30), indicating that the DNA substrate duplex remains intact at the PAM+2 site. Notably, beyond PAM+2, the dsMC95 DNA retains DNA-DNA pairing without engaging the RNA guide (Figure 2d, panel 3). Therefore, this configuration represents a “1-bp” R-loop state. In the alternative conformation (Figure 2c, panel 2b), dA66 of TS is flipped out of the DNA duplex and oriented toward the corresponding sgRNA uracil (rU19). This flipped out conformation appears to be stabilized by interactions between dA66 and a water molecule as well as by multiple contacts between the REC2 domain D269 side-chain and dA66, which have been reported in other structures with linear DNA substrates (e.g., PDB id 7S36^38^). dA66 and rU19 are separated by a distance of 4.4 Å, indicating a pre-hybridization geometry without stable pairing between the two nucleotides. Accordingly, this configuration is designated as a “1.5-bp” R-loop state that represents a transitional conformation between the stable 1-bp and 2-bp R-loop states. Notably, despite extensive 3D classification, these two configurations could not be further separated into distinct structural classes, indicating that State I captures an ensemble in which the 1-bp R-loop (PAM+2 DNA/DNA paired) and the 1.5-bp R-loop (PAM+2 DNA/DNA unwound) coexist in equilibrium.

**Table 1.**
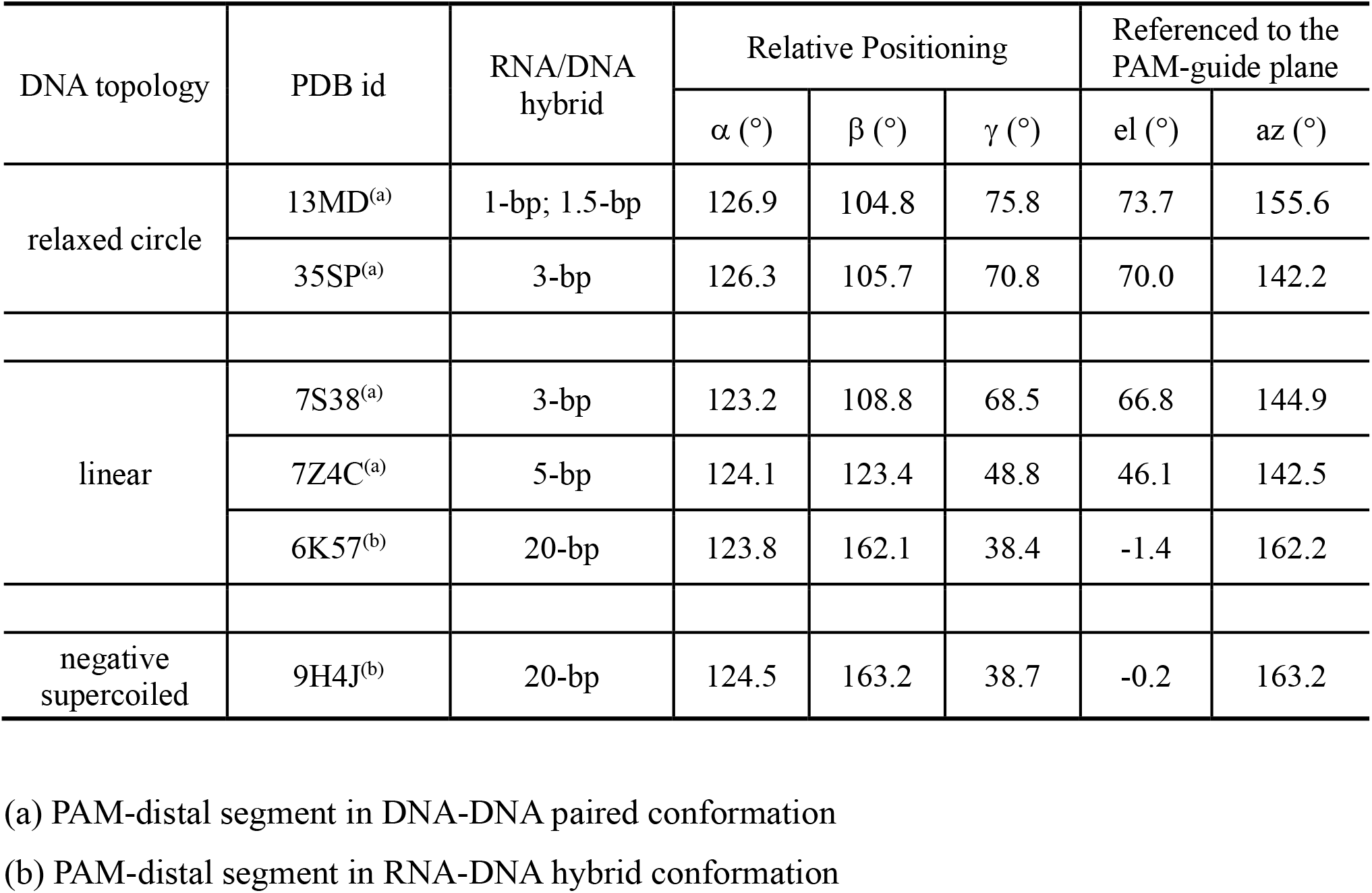
Spatial relationship between elements of PAM, RNA guide, and PAM-distal DNA.

| DNA topology | PDB id | RNA/DNA hybrid | Relative Positioning |  |  | Referenced to the PAM-guide plane |  |
| --- | --- | --- | --- | --- | --- | --- | --- |
| | | | $\alpha$ (°) | $\beta$ (°) | $\gamma$ (°) | el (°) | az (°) |
| relaxed circle | 13MD<sup>(a)</sup> | 1-bp; 1.5-bp | 126.9 | 104.8 | 75.8 | 73.7 | 155.6 |
|  | 35SP<sup>(a)</sup> | 3-bp | 126.3 | 105.7 | 70.8 | 70.0 | 142.2 |
| linear | 7S38<sup>(a)</sup> | 3-bp | 123.2 | 108.8 | 68.5 | 66.8 | 144.9 |
|  | 7Z4C<sup>(a)</sup> | 5-bp | 124.1 | 123.4 | 48.8 | 46.1 | 142.5 |
|  | 6K57<sup>(b)</sup> | 20-bp | 123.8 | 162.1 | 38.4 | -1.4 | 162.2 |
| negative supercoiled | 9H4J<sup>(b)</sup> | 20-bp | 124.5 | 163.2 | 38.7 | -0.2 | 163.2 |

(a) PAM-distal segment in DNA-DNA paired conformation
(b) PAM-distal segment in RNA-DNA hybrid conformation

Beyond the PAM+1 and PAM+2 positions, dsMC95 maintains DNA-DNA pairing, and the PAM-distal duplex segment makes contacts with the REC2 domain of dCas9 (Figure 3d). The positioning of the PAM-distal paired DNA duplex is identical to that reported in early R-loop intermediates formed with linear DNA substrates containing fewer than 2-bp of RNA/DNA hybrid. For example, a “0-bp” R-loop structure with a linear DNA substrate has been reported (PDB id 7S36,^38^ designated as “0-bp-linear”), in which the TS DNA PAM+1 and PAM+2 bases break pairing with the NTS DNA and flip toward the sgRNA guide without stable hybridization. Comparison of DNA between State I and this 0-bp-linear structure shows an overall RMSD of 0.5 Å (Figure S13), indicating very high similarity. Furthermore, in State I the Cas9 REC2 domain is positioned in close proximity to the DNA segment immediately beyond the R-loop, with N235 forming hydrogen bonds with the phosphate backbone of NTS PAM+9 (Figure 2d, panel 3). Such positioning of REC2 is similar to that observed in other early R-loop intermediate structures obtained with linear DNA substrates^38^ (Figure S13), and is consistent with a role of REC2 in stabilizing the early RNA/DNA hybrid (i.e., 1-bp and/or 2-bp R-loop) and maybe preparing the complex for further R-loop propagation.

**Figure 3.**
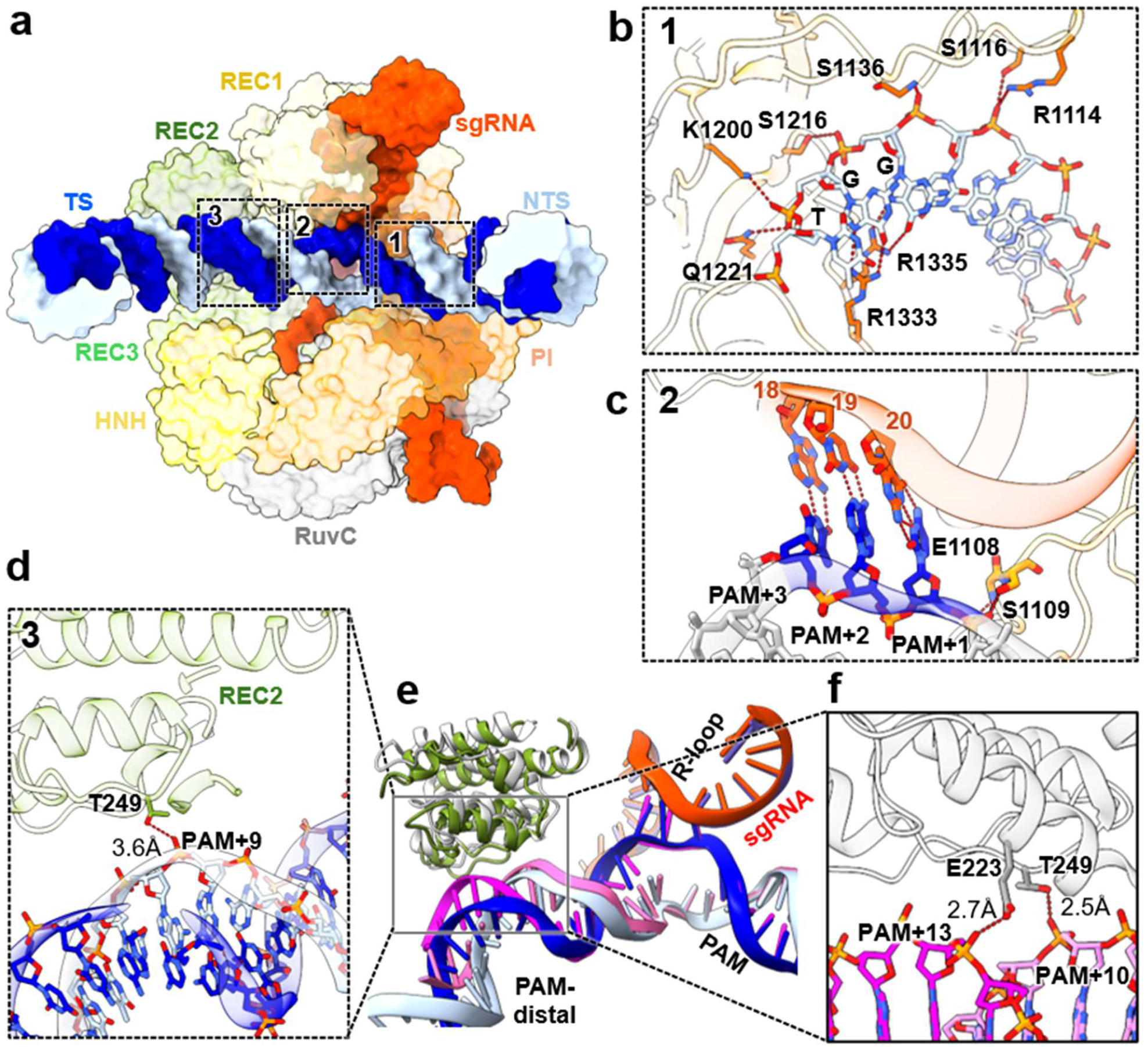
Molecular interactions within State II of the Cas9-dsMC95 complex. **a**, Surface representation of the Cas9-bound DNA segment. Cas9 domains are marked and color-coded as those in Figure 1b. The sgRNA is shown in orange, DNA TS in dark blue and NTS in light blue. Dashed boxes highlight the three key protein–DNA interaction regions with details shown in panels 1–3.. **b**, The PAM recognition interface, R1333 and R1335 within the PI domain directly engage the TGG PAM sequence through base-specific binding. Concurrently, the DNA backbone is stabilized by multiple amino acids, S1116, R1114, S1136, K1200, S1216, and Q1221. **c**, R-loop formation at the PAM-proximal region. A stable 3-bp R-loop spanning positions PAM+1 to PAM+3 is formed adjacent to the phosphate lock. **d**, Interaction between REC2 and the PAM-distal NTS. Residue T249 of the REC2 domain coordinates with the phosphate backbone of the NTS at the PAM+9 position with an interaction distance of 3.6 Å. **e**, Structural comparison with the linear DNA 3-bp R-loop structure (PDB: 7S38; Cas9 in gray, TS in pink, NTS in light pink). The two structures overlay nearly perfectly at the PAM, the sgRNA, and the 3-bp R-loop. With dsMC95, the paired PAM-distal DNA bends further away from the PAM and the REC2 domain shifts downwards, and the separation between the two increases as compared to that of the linear 3-bp R-loop structure. **f**, Close-up view of the corresponding region in the linear 3-bp R-loop structure. Residues T249 and E223 form close interactions with the DNA backbone at distances of 2.5 Å and 2.7 Å, respectively. These interactions appear to be weakened or absent in State II (panel **d**). See additional information in SI.

Overall, State I captures two new early R-loop states of Cas9 and reveals that the circular dsMC95 does not interfere with PAM recognition and R-loop initiation, but instead significantly affects R-loop propagation.

### State II reveals a native 3-base-pair R-loop state of Cas9 trapped by the circular dsMC95

Compared to State I, State II has a slightly better quality and higher resolution map, allowing building 95% of dCas9 residues (1,300 of 1,368 residues) including a majority of the HNH domain (Figure 3a; Figure S10). Similar to State I, 90% of the sgRNA (90 of 100 nucleotides) has been modeled, with the guide region (nucleotide 9–20) resolved to a local resolution of 2.5 Å (Figure S10a). The overall DNA map has a global resolution of 4.9 Å (Figure S8), however the PAM-10 to +10 segment of the dsMC95 was well resolved at ∼2 Å (Figure S10b&c). These features enabled the refinement of the complex including the entire 95-bp DNA circle (Figure S10c). The final model for State II has a MolProbity score of 1.25 (Table S2).

State II preserves several features observed in State I and other Cas9-bound DNAs. The sgRNA, including the single-stranded guide, adopts a highly similar conformation as that in the binary and other ternary complexes (Figure S14). The PAM recognition interface remains highly conserved, with PI domain residues R1333 and R1335 forming hydrogen bonding with the TGG PAM sequence (Figure 3b, panel 1; Figure S15a,b,c). Residues 1107-1109 form the phosphate lock with the DNA (Figure 3b, panel 1; Figure S15d,e,f), and the PAM+1 position shows a canonical RNA-DNA hybrid pair (Figure 3c, panel 2), indicating a successful R-loop initiation.

In contrast to State I, State II shows an extension of the R-loop to three complete hybrid pairs at the PAM+1 to +3 positions, with the RNA guide forming W-C pairing with the DNA TS, while the displaced NTS nucleotides remain clearly visible (Figure 3c, panel 2). Beyond the PAM+4 position, the DNA TS and NTS strands maintain pairing and bends away from the RNA guide (Figure 3d, panel 3). As such, State II captures a “3-bp” R-loop intermediate of Cas9. Furthermore, when State II is compared with other Cas9 R-loop structures, including that with a 3-bp RNA-DNA hybrid obtained using a linear DNA substrate with the two strands crosslinked (PDB id 7S38,^38^ designated as “3-bp-linear”), the PAM+1 to +3 RNA-DNA hybrid pairs are identical with an RMSD of 0.29 Å (Figure 3e). This indicates that the 3-bp R-loop structure captured in State II reports native interactions between Cas9 and DNA and is not biased by the circular constraint presented in dsMC95.

Nevertheless, differences are observed between State II and the linear 3-bp R-loop structure in the positioning of the PAM+4 to +14 segment of the protospacer, which maintain DNA/DNA pairing (Figure 3d,e,f). In State II, the PAM+4-14 DNA segment bends further away from the Cas9 RNP as compared to that in the 3-bp-linear structure (Figure 3e). Subsequently, although the REC2 domain in State II appears shifted toward the PAM-distal DNA, its stabilizing contacts with the DNA are diminished relative to the 3-bp-linear structure. For example, the hydrogen bond between E223 and the PAM+13 phosphate observed in the 3-bp-linear structure is lost, while the interaction between T249 and the DNA backbone is weakened (2.5 Å in 3-bp-linear versus 3.6 Å in State II) (Figure 3d,e,f). The more pronounced PAM-distal bending in State II likely results from the ring strain imposed by the closed minicircle, creating a substantial energetic barrier that hinders further R-loop propagation beyond the 3-bp stage. As a result, the topological constraints of the minicircle trap Cas9 in an early R-loop intermediate, preventing the large-scale conformational reengagements required for DNA cleavage.

### Early-stage R-loop propagation achieved by rigid-body swinging of the Cas9 RNP while dsMC95 maintains its global shape

Our State I and State II structures represent the only Cas9 ternary complexes in which the entire DNA substrate is fully resolved. This provides a unique opportunity to analyze Cas9-DNA interactions from both the perspective of the DNA and that of the Cas9 RNP. When the DNA circles from the 1.5-bp (State I) and 3-bp (State II) R-loop structures are superimposed, the RMSD across all 95 aligned pairs of the DNA is 3.1 Å (Figure 4a). However, upon excluding the 28-bp segment that encompasses the PAM and the protospacer, RMSD of the remaining 67 bp reduces to 1.1 Å (Figure 4a; Figure S16). This indicates that DNA structural variations are largely confined within the 1/3 of the ring interacting directly with Cas9, whereas the remaining 2/3 of the ring exhibit only minor structural differences. Furthermore, in both states, the overall DNA ring remains mostly flat and closely resembles the conformation of free dsMC95 (Figure 4a; Figure S17). As such, from the DNA perspective, dsMC95 maintains its global shape, deforming locally to accommodate early R-loop formation.

**Figure 4.**
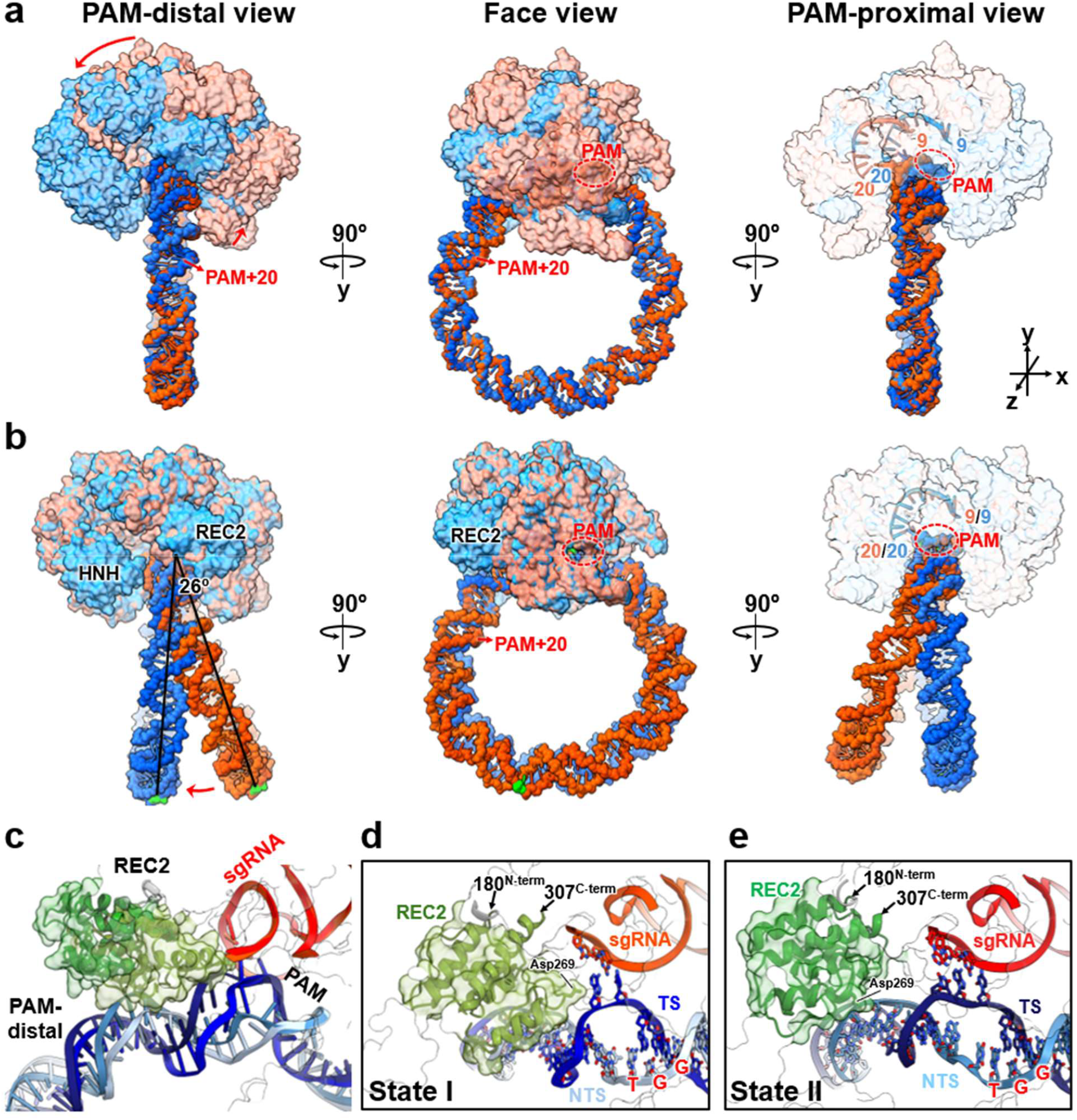
Comparisons between State I and State II of the Cas9-dsMC95 complexes. **a**, Overlay based on the DNA duplex spanning nucleotide 84-95-1-55 of the C-strand, which has an RMSD of 1.1 Å over the 67 pruned atom pairs. The State I complex shows the RNP in light orange and the DNA in orange. The State II complex shows the RNP in light blue and DNA in blue. In the PAM-distal view, the respective RNA guide spanning nucleotides 9–20 are shown as sticks. **b**, Overlay based on the RNP, which has an RMSD of 0.70 Å over 1,048 pruned Cα atom pairs. The rendering scheme is the same as that of panel **a**. The REC2 domain (marked) undergoes a distinct shift, whereas the HNH domain (marked) lacks resolved density in State I. The RNA guides are nicely aligned as shown in the PAM-proximal view. The black lines in the PAM-distal view mark, for the respective States I and II, the vector extending from the phosphate atom NTS dT64 (part of the PAM) and the phosphate atom of NTS dG14, which represents the plane of the DNA ring. A lateral swing of 26° is measured between the two vectors using ChimeraX. **c**, Closed-up view of the Cas9-bound DNA segment based on the RNP alignment. **d**, Close-up view of the REC2 domain in State I. REC2 is positioned close to the PAM+2 site, with Asp269 interacting with the flipped PAM+2 base. **e**, Close-up view of the REC2 domain in State II. Asp269 shifts away from the PAM+2 site and loses contact with the R-loop. The termini of REC2 are marked in panel **d** and **e** to aid comparison.

Superposition of dsMC95 in State I and State II reveals a rotation of the Cas9 RNP along the DNA (Figure 4a). To gain a perspective from the Cas9 enzyme, the two Cas9-dsMC95 complexes are superimposed based on the Cas9 protein (Figure 4b). The analysis yields an RMSD of 0.7 Å over 1,048 Cα atoms (1,048/1,205≈87% of the resolved Cas9 residues), with main differences observed at the REC2 domain and the poorly resolved HNH domain (Figure 4b). In addition, the sgRNA superimposes to each other with an RMSD of 0.63 Å (Figure 4b). Consistent with results obtained based on the DNA overlay (Figure 4a), the overall dsMC95 ring is highly similar in the XY-plane (Figure 4b, “Face view”), whereas a lateral swing of ∼26° is observed along the Z-direction (Figure 4b). Together, these observations indicate that the Cas9 RNP enzyme largely acts as a rigid body, attempting to induce “movement” of the dsMC95 ring to achieve early-stage R-loop formation.

The close alignment of the Cas9 RNP in State I and State II (Figure 4b) is consistent with comparison between the Cas9-dsMC95 complexes and Cas9-bound linear DNAs at a comparable R-loop stage (Figures 2, 3), which shows a very high degree of structure similarity in sgRNA engagement, PAM interaction, and R-loop initiation. Furthermore, when aligned based on the Cas9 protein, the PAM+1-3 RNA-DNA hybrid pairs also overlay nearly completely between the 3-bp-linear R-loop structure and State II, which contains the circular dsMC95 (Figure 3e). These results indicate that during the process of engaging a DNA substrate to establish the early-stage R-loop, the Cas9 RNP enzyme maintains the same “rigid body” configuration regardless of the overall topology of the DNA substrate.

Nevertheless, the DNA ring “shape” does manifest itself. When the Cas9 protein is superimposed, both the RNA guide and the PAM overlay well, but differences are observed between the DNA PAM-distal protospacer segment (i.e., PAM+4 to +14), which is right next to the RNA-DNA hybrid and maintains DNA-DNA pairing (Figure 4c). As the R-loop extends to 3-bp, the PAM-distal paired DNA shifts away from the PAM and moves closer to the pre-set RNA guide (Figure 4c). Accompanying the change in the PAM-distal DNA positioning, the REC2 domain (residues 177–305) also undergoes a clear conformational shift (Figure 4c). In the 1.5-bp R-loop structure (State I), REC2 is positioned proximally to both the RNA-DNA hybrid and the PAM-distal DNA duplex, with Asp269 contacting the flipped TS PAM+2 base (Figure. 4d). On the other hand, in the 3-bp R-loop structure (State II), REC2 moves away from the RNA-DNA hybrid and the PAM-distal paired DNA (Figure 4e). Interestingly, in the “3-bp-linear” R-loop structure (PDB id 7S38^38^), REC2 moves away from the RNA-DNA hybrid pairs, however, the paired PAM-distal DNA duplex also moves to remain close to REC2 (Figure 3f). Together these observations suggest that REC2 and PAM-distal DNA move in coordination after forming a stable 3-bp R-loop.

### Interplay between ring constraints and DNA unwinding impact R-loop propagation

The analysis described above reveals that a key distinguishing feature among the Cas9-bound DNA structures is the positioning of the PAM-distal paired DNA-DNA segment. To further quantify the movement of the paired PAM-distal DNA, we analyzed the relative orientations of three vectors representing the PAM DNA duplex (designated as the “PAM” vector), the sgRNA guide (“guide” vector), and the PAM-distal paired DNA duplex (“distal” vector), respectively (Figure 5a). Consistent with Cas9-based superimposition (Fig. 4b, 4c), between the 1.5-bp and 3-bp R-loop structures, the angle between the PAM and guide vectors (α, Figure 5a) remains nearly the same (126.9° and 126.3°, respectively, Table 1), reflecting invariant positioning between the RNA guide and the DNA PAM. On the other hand, the angle between the PAM vector and the distal vector (β, Figure 5a) increases (104.8° and 105.7°, respectively), while the angle between the guide vector and the distal vector (γ, Figure 5a) decreases (75.8° and 70.8°, respectively, Table 1), indicating that the PAM-distal duplex moves away from the PAM while positions closer to the RNA guide. Furthermore, with respect to the plane defined by the PAM vector and the guide vector, the elevation (el) and azimuth (az) angles of the distal vector (Fig. 5a) are 155.6° and 73.7°, respectively for the 1.5-bp R-loop and 142.2° and 70.0° for the 3-bp R-loop (Table 2). These indicate the PAM-distal DNA undergoes a combined motion of twisting (rotation) toward PAM-guide plane and swinging toward the guide within the PAM-guide plane.

**Figure 5.**
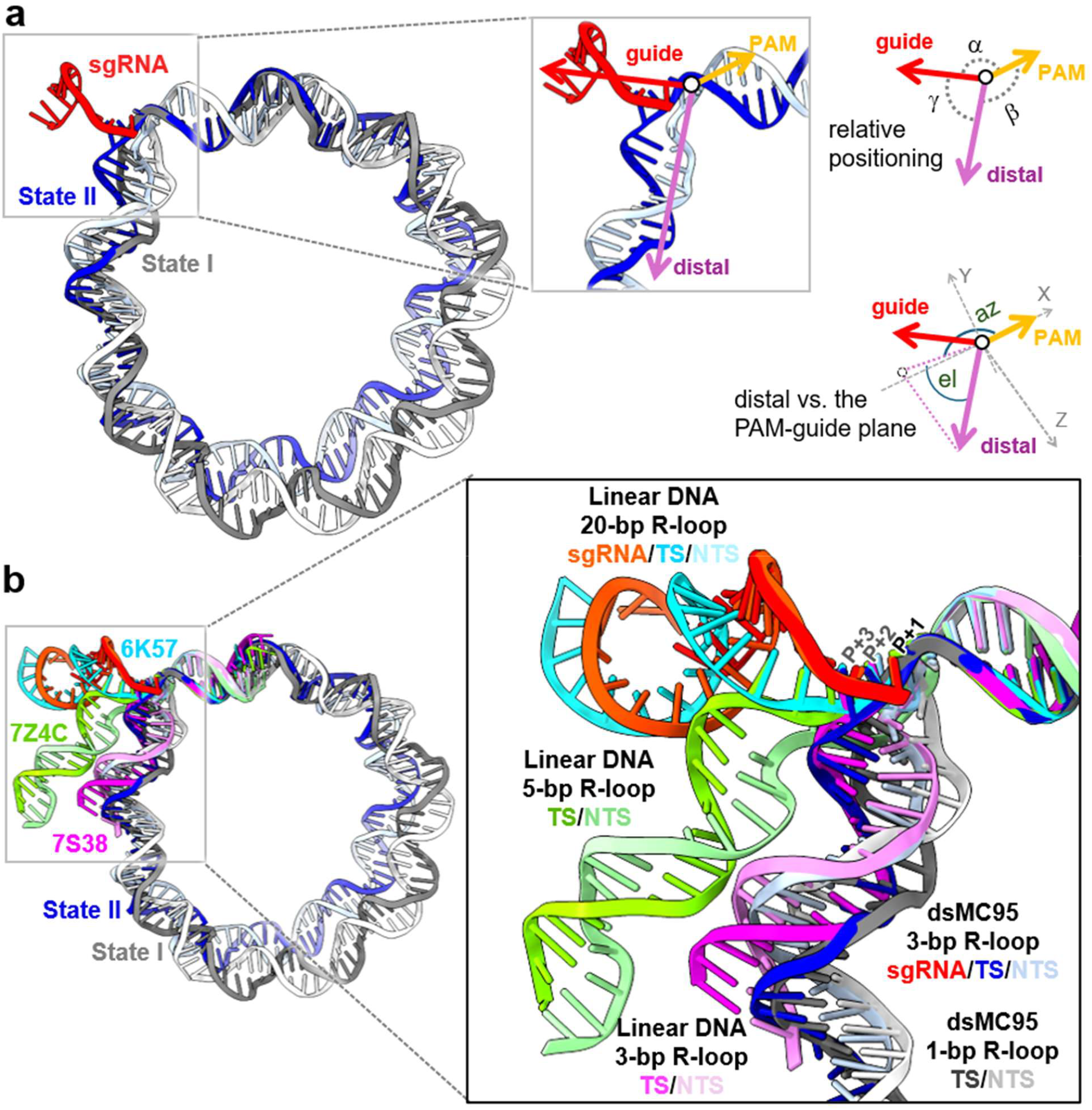
Correlating PAM-distal DNA movement with R-loop propagation. **a**, Analysis of PAM-distal DNA positioning. Shown on the right is overlay of dsMC95 State I (gray) and State II (dark blue) based on the RNP alignment, which shows deviation between the DNA rings. The inset illustrates the PAM vector (yellow), the guide vector (red), and the distal vector (magenta), which represent, respectively, the PAM, the RNA guide, and the PAM-distal DNA (see Methods). Shown on the left are illustrations of the angles characterizing relative orientation of the three vectors (top) and the positioning of the PAM-distal DNA with respect to the plane defined by the PAM and guide vectors (bottom). **b**, Comparisons between Cas9-bound dsMC95 with DNA observed in ternary complexes obtained with linear duplexes and adopting various R-loop states. The full DNA view is shown on the right, and an enlarged view focusing on the PAM-distal DNA is shown on the left.

Further analysis of the relative orientations between the PAM, guide, and PAM-distal vectors were carried out with other Cas9 ternary complexes (Figure 5b, Table 1). With linear DNA substrates, as the R-loop progress from 3-bp (PDB id 7S38^38^) to 5-bp (PDB id 7Z4C^12^) and 20-bp (PDB id 6K57^39^), α remains nearly identical, reflecting the invariant positioning of the RNA guide and the PAM. β increases monotonically, indicating that the PAM-distal DNA bends away from the PAM; while at the same time, γ decreases monotonically, reflecting that the PAM-distal DNA is pulled toward the pre-positioned RNA guide. In addition, with respect to the PAM-guide plane, the distal vector shows monotonically decreasing elevation. This indicates that the PAM-distal paired DNA progressively swings toward the pre-positioned RNA guide as the R-loop propagates,^38,12,13^, which is consistent with conformational changes observed in Cas9-bound dsMC95 structures (Figure 5a, Table 1).

Furthermore, structures of Cas9 bound to negatively supercoiled (-sc) DNA minicircles have recently been reported.^31^ Analysis of one of these structures (PDB id 9H4J^31^), featuring a stable 20-bp R-loop, shows that the relative orientation between the PAM, guide, and distal vectors is nearly identical to that found with the “20-bp-linear” R-loop structure (PDB id 6K57) (Table 1; Figure S18). Together, these data suggest that the tilt-and-swing motion of the PAM-distal paired DNA is a common mechanism used by Cas9 to propagate the R-loop, regardless of the DNA topology.

While Cas9 utilizes the same general mechanism to unwind target DNA and propagate the R-loop, the manner in which the DNA (in particular the PAM-distal paired segment) responds to Cas9 actions ultimately determines whether R-loop propagation proceeds successfully (Figure 5). With the dsMC95 minicircle, the PAM-distal DNA experiences a tug-of-war between two opposing forces. As the action of Cas9 drives the DNA away from PAM to meet the pre-positioned RNA guide, the constraints of the closed-ring topology try to keep the DNA bend and stay away from the RNA guide (Figure 5). As a result, the protospacer cannot be efficiently unwound to form a stable R-loop, which in turn weakens Cas9 binding (Figure S3) and impairs DNA cleavage (Figure S5).

It is intriguing that dsMC95, a relaxed 95-bp minicircle, stalls the R-loop propagation in an early stage; while the (-sc) 126-bp minicircle supports formation of a stable full R-loop even in the presence of RNA-DNA mismatches.^31^ Structural comparisons between Cas9-bound dsMC95 and (-sc) 126-bp minicircle (9H4J^31^) again show nearly identical RNA-guide and PAM positioning, while the DNA trajectories diverge (Figure S18). However, with the (-sc) 126-bp minicircle, a large portion of the DNA ring is unresolved in the Cas9 complex,^31^ rendering it difficult to compare it with dsMC95. In the future, more in-depth studies of the interactions between DNA minicircles and Cas9 will be of considerable interest.

## Conclusion

The high-resolution cryo-EM structures reported here provide important insights into the interplay between DNA shape and Cas9 target recognition. Regardless of DNA topology, Cas9 appears to function largely as a rigid-body during PAM searching, R-loop initiation, and R-loop propagation. These Cas9 actions not only cause unwinding of the DNA protospacer duplex, but also require DNA swinging motion toward the pre-positioned RNA-guide. The relaxed dsMC95 minicircle possesses sufficient flexibility to deform and accommodate the early R-loop states, while its close-ring topology constrains the PAM-distal paired DNA movements, thereby interfering with further R-loop propagation and DNA cleavage. These data, therefore, unambiguously demonstrate that conformational dynamics of Cas9, and consequently its function, are affected by intrinsic DNA shape features, in this case the relaxed circle topology, resulted from sequences extending well beyond the PAM and the protospacer. In the complex genome environment, variation in DNA topology (e.g. supercoiling) as well as binding by other proteins (e.g., nucleosome) are likely to affect such PAM-distal movements, thus modulating Cas9 binding and cleavage. Further studies are needed to elucidate the relationship between DNA global shape features and Cas9 function, as well as to leverage these insights to guide Cas9 target selection.

## Acknowledgements

Electron microscopy data were collected at the Core Center of Excellence in Nano Imaging (CNI) at University of Southern California (USC). Cryo-EM data processing was carried out at the Center for Advanced Research Computing (CARC) at USC. We thank Dr. H. Khant (USC) for assisting with cryo-EM data acquisition, and Dr. Tomek Osinski (USC) for assisting with cryo-EM data processing.

## Author contributions

Conceptualization, KY.L., Y.L., and P.Z.Q.; Methodology, KY.L., D.K., Y.L., J.J., and P.Z.Q.; Investigation, KY.L., D.K., Y.L., H.C. J.J., V.C.; Formal Analysis, KY.L., D.K., J.J., V.C., and P.Z.Q; Writing – Original Draft, KY.L. and D.K.; Writing – Review & Editing, KY.L., D.K., J.J., V.C., and P.Z.Q. All authors reviewed the manuscript.

## Supplementary Data

Supplementary Information is available online.

## Data Availability

The atomic models have been deposited in the Protein Data Bank (PDB) under accession codes 13MD for State I and 35SP for State II. The cryo-EM maps have been deposited in the Electron Microscopy Data Bank (EMDB) under accession codes EMD-77151 for the composite map of State I, EMD-77147 for the consensus map of State I, EMD-77150 for the dsMC95-focused map of State I, EMD-77167 for the composite map of State II, EMD-77165 for the consensus map of State II, and EMD-77166 for the dsMC95-focused map of State II.

## Materials and Methods

### Expression and purification of Cas9

The *Streptococcus pyogenes* catalytically inactive Cas9 (dCas9, Addgene pMJ841, with D10A/H840A mutations) protein with an N-terminal His_6_-MBP-TEV fusion tag to facilitate purification was expressed in *E. coli* Rosetta2 (DE3) pLysS (Novagen) for 16 hours at 18 °C. Bacterial cells were harvested and lysed in a buffer containing 20 mM HEPES pH 7.0, 500 mM NaCl, and 5% glycerol. Following centrifugation to clarify the lysates, the supernatant was applied to a 2 ml TALON Superflow IMAC column (Cytiva), washed with 10 column volumes of the same lysis buffer supplemented with 5 mM imidazole. Tagged Cas9 was eluted with 5 column volumes of 20 mM HEPES pH 7.0, 250 mM NaCl, 5% glycerol, and 200 mM imidazole. The His_6_-MBP tag was removed by overnight cleavage with TEV protease at 4 °C. The salt concentration was then decreased to 150 mM NaCl by dilution, and the sample was applied to a 1 ml HiTrap SP HP cation exchange column (Cytiva) equilibrated in 20 mM HEPES pH 7.0. After washing with buffer containing 150 mM NaCl, Cas9 was eluted over a linear gradient of 0–100% high salt buffer (up to 1.0 M NaCl), with peak elution around 400 mM NaCl. The protein was concentrated and further purified by size-exclusion chromatography using a Superdex 200 Increase 10/300 GL column (GE Healthcare) in 20 mM HEPES pH 7.0, 150 mM NaCl, and 0.5 mM DTT. The final purified protein fractions were concentrated to 5 mg/ml and stored at -80 °C. The Wide-type Cas9 (WTCas9, Addgene pMJ806) used in the assay was purified using the same protocol as described above.

### *In vitro* sgRNA transcription

The single-guide RNA (sgRNA) was synthesized by T7 *in vitro* transcription a following reported procedure.^32^ using a 400 µL transcription reaction containing 40 mM Tris pH 7.5, 15 mM MgCl_2_, 2 mM spermidine, 0.01% Triton X-100, 1 mM each of CTP, ATP, GTP, and UTP, 5 mM DTT, 7.5 nM DNA template, and 30 µg of T7 RNA polymerase. The reaction mixture was incubated at 37 °C for 4 hours. Following transcription, sgRNA was precipitated with ethanol overnight, purified by electrophoresis on a 10% denaturing polyacrylamide gel containing 7 M urea, subjected to a second round of ethanol precipitation, and resuspended in nuclease-free water. The sequence of sgRNA is detailed in Supplementary Table 1.

### Synthesis of the 95-basepair DNA minicircle

Equimolar amounts of 5′-phosphorylated oligonucleotide hairpins A1 and A2 were combined in 70 μL of TE buffer (pH 8.0) to a final concentration of 3.5 μM per strand. The mixture was annealed by slow cooling from 95 °C to room temperature over 3 hours. Subsequently, 4 μL of T4 DNA ligase (5 Weiss units/μL) in ligation buffer containing 66 mM Tris-HCl pH 7.6, 6.6 mM MgCl_2_, 10 mM DTT, 0.1 mM ATP was added, and the reaction was incubated at 16 °C for 16 hours. T4 ligase was inactivated by heating the reaction to 65 °C for 10 min. The resulting DNA dumbbell (A strand) was purified by electrophoresis on a 10% denaturing polyacrylamide gel, followed by ethanol precipitation. For the assembly of the 95-bp double-stranded minicircle (dsMC95), equimolar amounts of the purified DNA dumbbell A strand and two 5′-phosphorylated hairpins B1 and B2 were mixed to a final concentration of 1.5 μM for each component. Annealing was performed by gradually cooling from 95 °C to room temperature over the course of one night. Ligation was carried out using the same conditions as described for strand A. The final dsMC95 product was purified by 10% denaturing PAGE and ethanol precipitation. The sequences of A1, A2, B1, B2, and the final dsMC95 are provided in Table S1.

### Assembly and purification of the dCas9-sgRNA-dsMC95 complexes

The dsMC95-bound Cas9-sgRNA complex was assembled in a reaction buffer containing 20 mM Tris pH 7.5, 100 mM KCl, and 5 mM MgCl_2_, using a 1:1.5:1 molar ratio of Cas9:sgRNA:dsMC95. The 20-nt sgRNA was denatured at 95 °C for 1 min and then allowed to cool to room temperature. Cas9 and sgRNA were incubated together for 10 min at room temperature to form the binary complex before the addition of the 95-bp dsMC95 substrate. The reaction mixture was incubated at 37 °C for 45 min to form the ternary Cas9-sgRNA-dsMC95 complex. The resulting complex was concentrated and injected into a Superdex 200 Increase 10/300 GL column (GE Healthcare) equilibrated in the same buffer. Final samples were concentrated to 5 mg/mL for cryo-EM using an Amicon Ultra Centrifugal Filter (50 kDa molecular weight cutoff).

### Cryo-EM grid preparation and data collection

Cryo-EM grids were prepared by applying 3 µL of Cas9-sgRNA-dsMC95 samples onto glow-discharged Quantifoil holey carbon grids (Au, R1.2/1.3, 300 mesh) and blotting for 3 s, followed by plunge-freezing into liquid ethane using a Vitrobot Mark IV (Thermo Scientific) operating at 100% humidity and 4 ºC. Cryo-EM data were collected using a Krios microscope operating at 300 kV with a K3 direct electron detector at a nominal magnification of 105,000x resulting in the pixel size of 0.85 Å. 7,503 movies were recorded in 50 frames with a defocus range of -1.0 to -2.4 µm and a total dose of 51.3 e^-^/A^2^.

### Image processing and 3D map construction

Data processing was performed using cryoSPARC (v4.7.1)^40^ unless stated otherwise. Initial alignment of cryo-EM movie stacks was carried out using Patch motion correction, followed by Patch CTF estimation. Micrographs were curated based on a CTF resolution cut off less than 4.0 Å, resulting in 5,091 corrected micrographs selected for analysis. Particle picking was initially conducted using the blob-picker, followed by particle extraction (box size: 400 pixels, Fourier-cropped to 64 pixels, with a pixel size of 5.3125 Å/pix) and multiple rounds of 2D classification. The best particles from this step were then used as templates for template-based particle picking. For the Cas9-sgRNA-dsMC95 complexes, a total of 4,621,541 particles were picked and extracted (box size of 400 pix and Fourier cropped to 64 pix, 5.3125 Å/pix) followed by two rounds of iterative 2D classification. From this, 1,269,686 particles displaying features consistent with a single Cas9-sgRNA complex bound to one dsMC95 were selected, and 102,823 particles from 1,000 movies were used for Topaz train, followed by Topaz extraction and particle extraction (box size of 400 pix and Fourier cropped to 64 pix, 5.3125 Å/pix), which produced 1,334,414 particles. The duplicates between the particles selected from template picking and Topaz picking were removed and followed by *Ab initio* and 4× iterative heterogeneous refinements. The cleaned particle stack of the selected 3D class underwent 2^nd^-round *Ab-initio* and heterogeneous refinement. The final 49,515 and 313,596 particles for Class I and Class II of the dCas9-sgRNA-dsMC95 complexes, respectively, were extracted in full resolution (box size of 400 pix, 0.85 Å/pix), and additional 3D classification was performed for Class I to clean up the junk particles. Non-uniform refinement resulted in gold-standard Fourier shell correlation (GSFSC) resolutions of 3.08 Å and 2.67 Å Class I and Class II, respectively. To clearly observe dcMC95, protein signal was removed using the dCas9-focused mask from the cleaned particle stack by particle subtraction tool, and the resulting particle stacks were locally refined using the dsMC95-focused mask, producing 6.59 Å and 4.87 Å maps of dsMC95 for Class I and Class II, respectively. The final composite maps were generated by combining consensus maps and dsMC95-focused maps using the Combine Focused

Maps tool in Phenix^41^ at an overall nominal resolution of 3.1 Å and 2.7 Å using the FSC=0.143 criterion for Class I and Class II, respectively. The final maps were sharpened using the sharpening tool and their local resolution was estimated in cryoSPARC (v4.7.1).^40^

### Model building and refinement

The initial SpyCas9-sgRNA-DNA model (PDB ID 6K57)^39^ was fit into the obtained cryo-EM density map by ChimeraX.^42^ Subsequent refinement involved multiple rounds of real-space refinement with Phenix,^41^ along with manual model adjustment and validation in COOT.^43^ Structural figures were generated using ChimeraX^42^ and Pymol (Schrödinger). Data collection and refinement statistics are summarized in Table S2.

### Analysis of relative positioning of PAM-distal DNA

To quantify positioning of the PAM-distal DNA in a given Cas9 ternary complex, the average of C1’ coordinates of TS strand dA64 and NTS strand dT32, which represents the mid-point of the “dA/dT” pair of the “TGG” PAM (Figure S1), was chosen as the origin for all vectors. The PAM vector 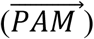 was defined as ending at the point represented by the average of C1’ coordinates of TS strand dC62 and NTS strand dG34 (Figure S1). The guide vector 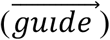 was defined as ending at the C1’ of the rC15 nucleotide (Table S1). In structures where the PAM-distal DNA-DNA pairing is maintained, the distal vector 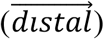 was defined as ending at the mid-point of the DNA-DNA base pair at the PAM+10 position, computed as the average of C1’ coordinates of the corresponding nucleotides. In structures where the PAM-distal DNA hybridizes with the RNA guide, 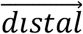 was defined as ending at the mid-point of the RNA-DNA hybrid pair at the PAM+10 position, computed as the average of C1’ coordinates of the corresponding nucleotides. The angle α between the PAM and the guide vectors was computed based on the dot products:

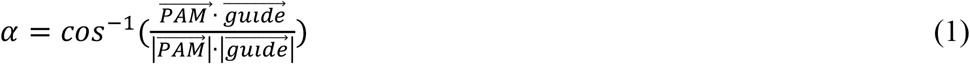

The angle β between the guide and the distal vectors was computed based on the dot products:

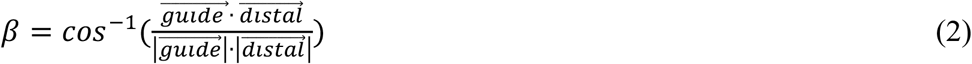

The angle γ between the PAM and the distal vectors was computed based on the dot products:

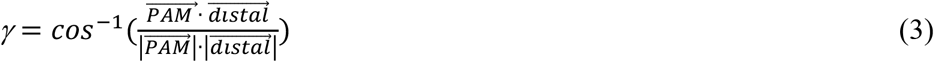

Furthermore, a new coordinate system, designated as the PAM coordinate, was defined with the origin of the coordinate system setting at the origin of the three vectors. The 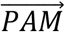 vector set as the X-axis 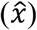:

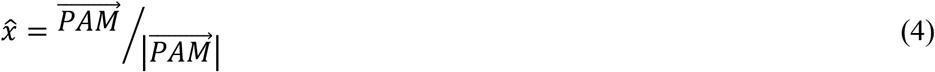

the XY-plane was defined the by 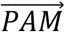 and the 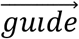 vectors, so that the Z-axis of the PAM coordinate 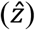 was calculated as the cross product of 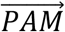 and 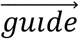:

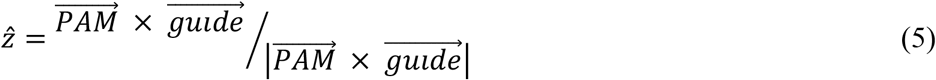

the Y-axis of the PAM coordinate *ŷ* was then calculated as the cross product of 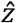 and 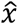:

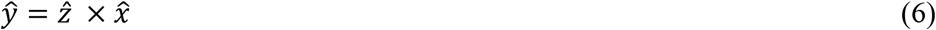

The elevation angle (el) of the distal vector with respect to the PAM-guide plane (i.e., the XY-plane) was computed from its Z-component expressed in the PAM coordinate:

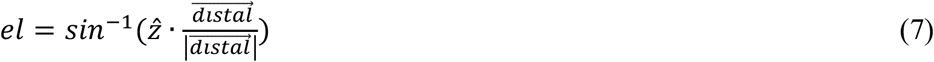

values of *el* span a range of (–90°, +90°), with 0° to +90° corresponding to 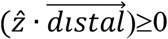, and – 90° to 0° corresponding to 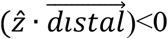.

The azimuth angle (az) of the distal vector with respect to the PAM-guide plane was computed from its X-and Y-components in the PAM coordinate:

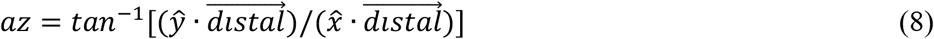

values of *az* span a range of (–180°, +180°), with 0° to +90° corresponding to 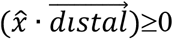 and 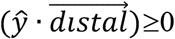; +90° to +180° corresponding to 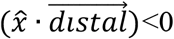 and 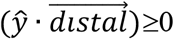 to 0° corresponding to 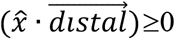 and 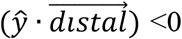; and –180° to –90° corresponding to 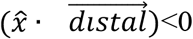 and 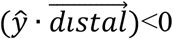;

