## Supplementary Information for "Structures of Cas9-Bound Double-Stranded DNA Mini-Circle Reveal Impacts of DNA Shape on Cas9 Target Interrogation"

**Supplementary Figure S1**

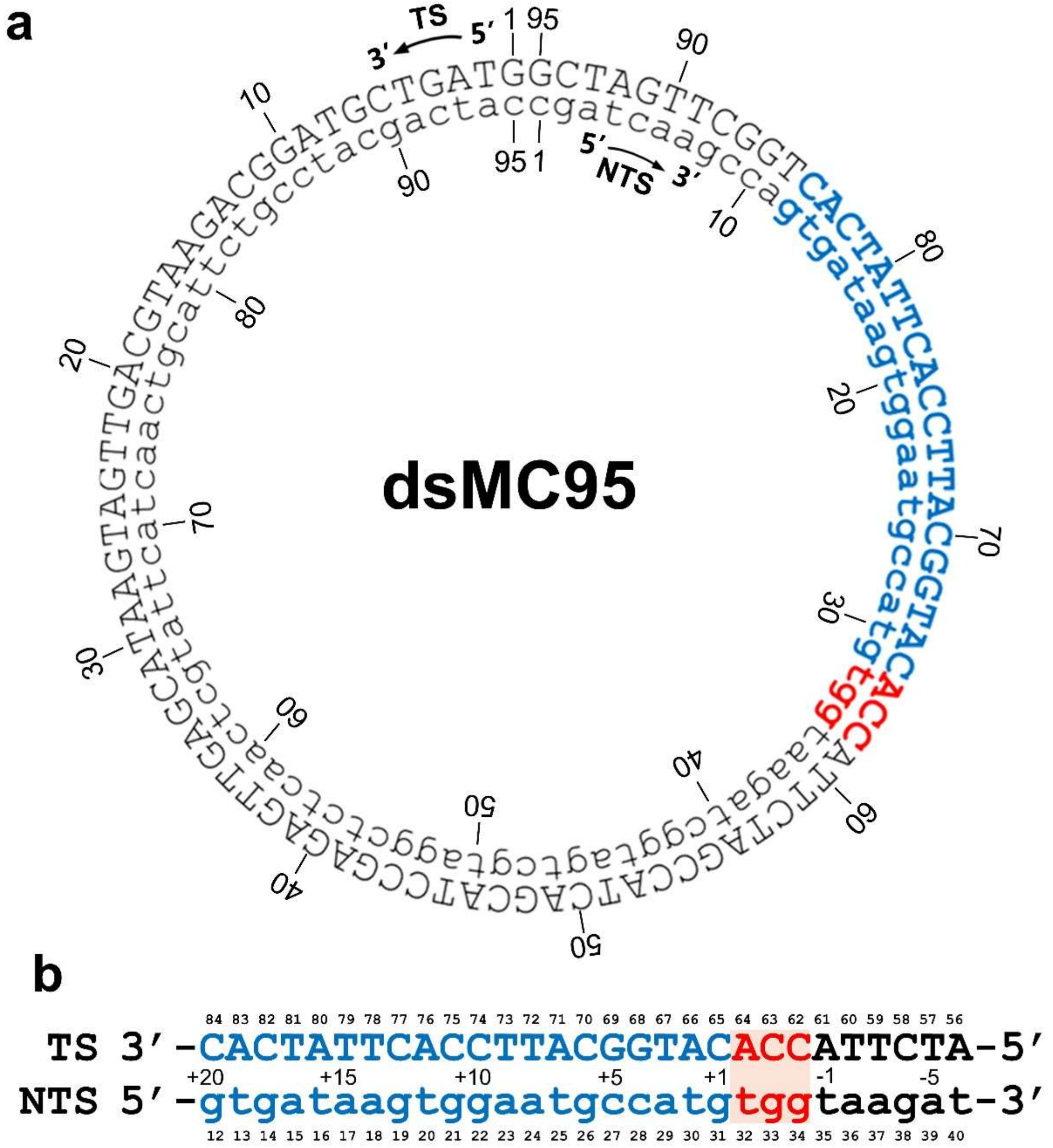


**Figure S1. Sequence and organization of the 95-basepair DNA minicircle (dsMC95). a**, Sequence of dsMC95 shown in a circular form. Following nomenclature used in Cas9 studies, the two strands are designated as the target strand (TS, uppercase) and the non-target strand (NTS, lowercase). The protospacer sequence is highlighted in blue, and the protospacer-adjacent motif (PAM) in red. Strand orientation is indicated by 5′ and 3′ labels. Nucleotide positions are marked at 10-bp intervals. dsMC95 was assembled from four synthetic hairpins (see Methods), with hairpins A1 and A2 corresponding to NTS sequence, whereas hairpins B1 and B2 corresponding to TS sequence (see Table S1). Although the circular DNA has no terminus, nucleotides are numbered from 1 to 95 based on the hairpin sequences used for assembly. The same numbering scheme is applied throughout the structural analysis and in the deposited PDB coordinates. **b**, Magnified view of the protospacer and protospacer-adjacent motif (PAM) regions. The nucleotide numbering relative to the PAM position (PAM-6 to PAM+20) is indicated, with the PAM sequence highlighted in red.

**Supplementary Figure S2**

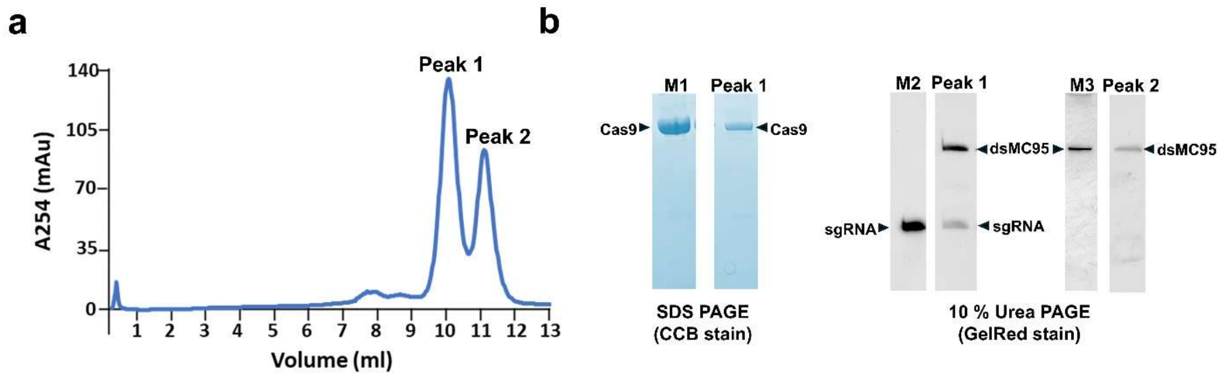


**Figure S2. Purification of the dCas9–sgRNA–dsMC95 ternary complex. a**, Representative size-exclusion chromatogram of the dCas9–sgRNA complex incubated with the 95-bp DNA minicircle (dsMC95). **b**, Peak validation by PAGE analysis. SDS-PAGE with Coomassie Brilliant Blue (CBB) staining shows the presence of Cas9 in Peak 1. Denaturing Urea PAGE with GelRed staining identifies both dsMC95 and sgRNA in Peak 1 (ternary complex), whereas Peak 2 contains only free dsMC95. Fractions corresponding to Peak 1 were pooled and used for cryo-EM grid preparation.

### Supplementary Figure S3

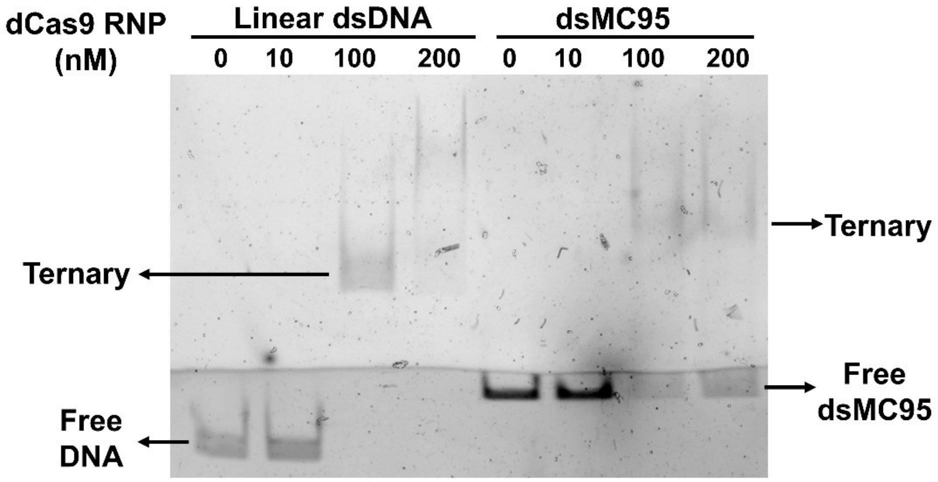


**Figure S3. dCas9 binding with dsMC95.** Electrophoretic mobility shift assay (EMSA) was used to examine binding between dCas9 RNP and DNA. As shown 10 nM DNA was incubated with the indicated concentration of dCas9 RNP in the reaction buffer (20 mM Tris pH 7.5, 100 mM KCl, and 5 mM $MgCl_2$). The mixture was incubated at 37 °C for 30 min, then run on a stacking (5% upper; 8% lower) native polyacrylamide gel. Samples were visualized by fluorescent imaging. The linear duplex consisted of the A1-40-FAM and B1 strands (Table S1). At 100 and 200 nM RNP, the linear duplex shifted nearly completely into the slow migrating band(s) representing the ternary complex, while a sizable amount of free dsMC95 remained. This indicates that the circular dsMC95 binds weaker to dCas9.

**Supplementary Figure S4**

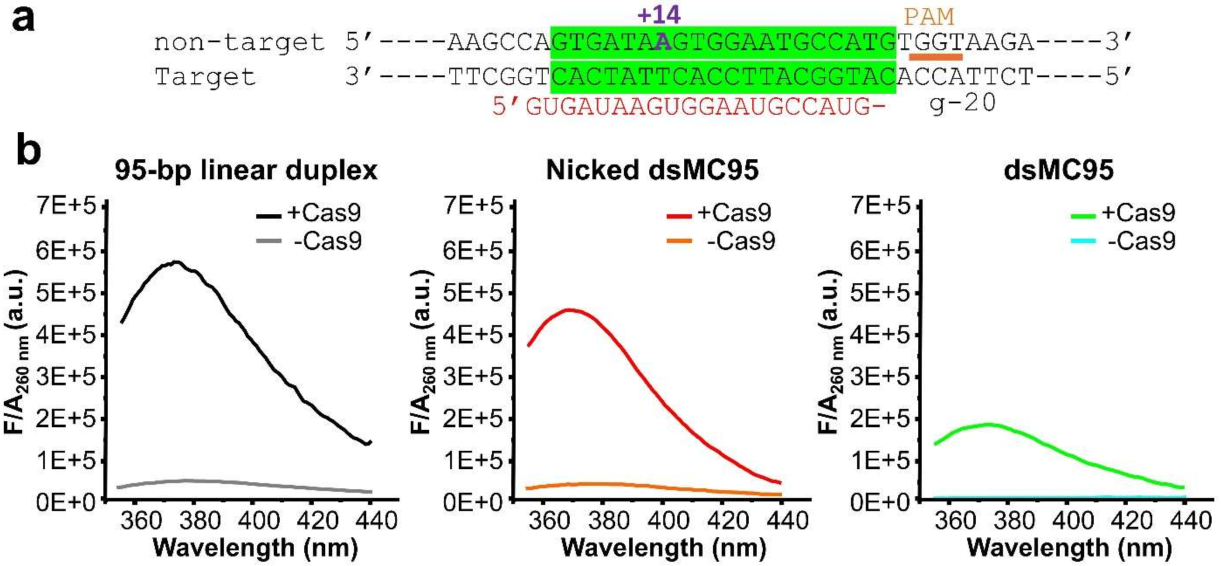


**Figure S4. Characterization of dCas9-induced DNA unwinding using 2-aminopurine (2AP) fluorescence. a**, DNA constructs used for 2AP fluorescence measurements. The DNA segment embedded in the measured constructs is shown, with the protospacer (green box) and the PAM (orange underline) marked. 2AP is substituted at the PAM+14 position of the non-target-strand. **b**, Representative 2AP fluorescence emission spectra of duplexed DNA with the same sequence but in different topological forms, a 95-bp linear duplex (left), nicked dsMC95 (middle), and covalently closed dsMC95 (right). 2AP-substituted DNA samples were prepared using the A1-14-2AP strand listed in Table S1, and ternary complex with dCas9 RNP was formed as described in Figure S3 and purified by size-exclusion chromatography (Figure S2). 2AP fluorescence emission measurements were carried out using a SpectraMax iD5 microplate reader (Molecular Devices, San Jose, CA), with each sample containing approximately 1 μM DNA. Excitation was set at 320 nm, and temperature was maintained at 25 °C during data acquisition. Absorbance was obtained immediately after the fluorescence measurement on a LAMBDA UV/Vis/NIR Spectrophotometer (Perkin-Elmer). The background-corrected emission spectrum (F, obtained by subtracting the corresponding buffer emission) was normalized by $A_{260}$ of the same sample. The resulting $F/A_{260}$ values, which are proportional to the fluorescence quantum yield of 2AP, was used to assess DNA base stacking. With the closed dsMC95 (right), the presence of dCas9 RNP ("+Cas9") increases $F/A_{260}$, indicating reduced DNA base stacking due to dCas9 unwinding of the protospacer at position PAM+14. However, $F/A_{260}$ increase with the closed dsMC95 (right) is smaller compared to that of the nicked circle (middle) and linear duplex (left). This likely reflects reduced binding of dsMC95 (Figure S3) as well as reduced unwinding in the bound dsMC95.

**Supplementary Figure S5**

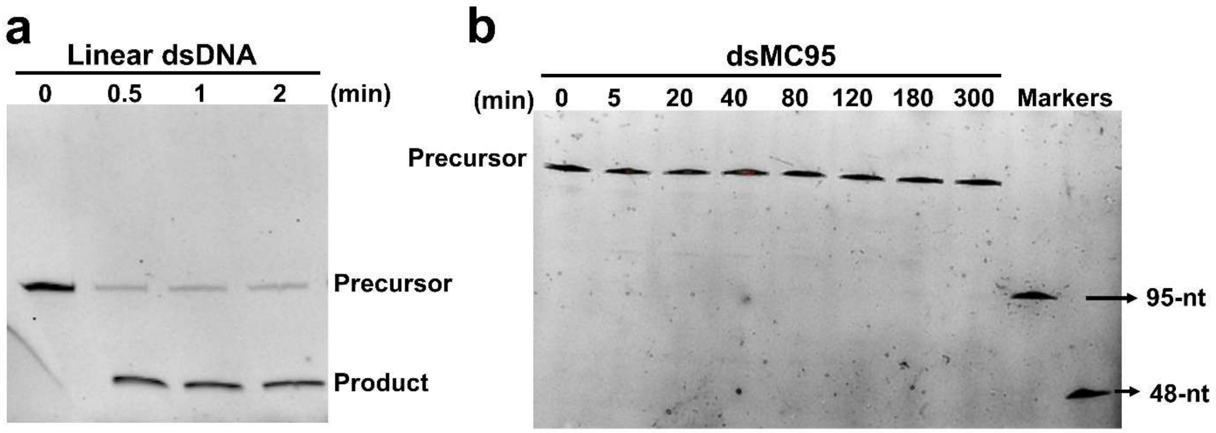


**Figure S5. Assessing Cas9 cleavage of dsMC95**. Cleavage assay was carried out at 37°C in the reaction buffer (Figure S2) using 10 nM DNA substrate, 600 nM SpyCas9, and 720 nM g-20 sgRNA (Table S1). Samples were resolved on a 10% denaturing polyacrylamide gel and visualized via fluorescence. **a**, Cas9 cleavage with a linear DNA duplex assembled using the A1-40-FAM and B1 strands (Table S1). The DNA precursor was converted to the expected product within 0.5 min, indicating that the Cas9 RNP is fully active. **b**, Cleavage of dsMC95 conducted in parallel with that of linear duplex. The markers are respectively a 95-nt linearized DNA (corresponding to TS in Table S1) and the 48-nt A1-40-FAM strand (Table S1). No detectable time-dependent product generation was observed at 300 min nor with extending the incubation time to 12 hr (data not shown). The results indicate no dsMC95 cleavage by Cas9.

**Supplementary Figure S6**

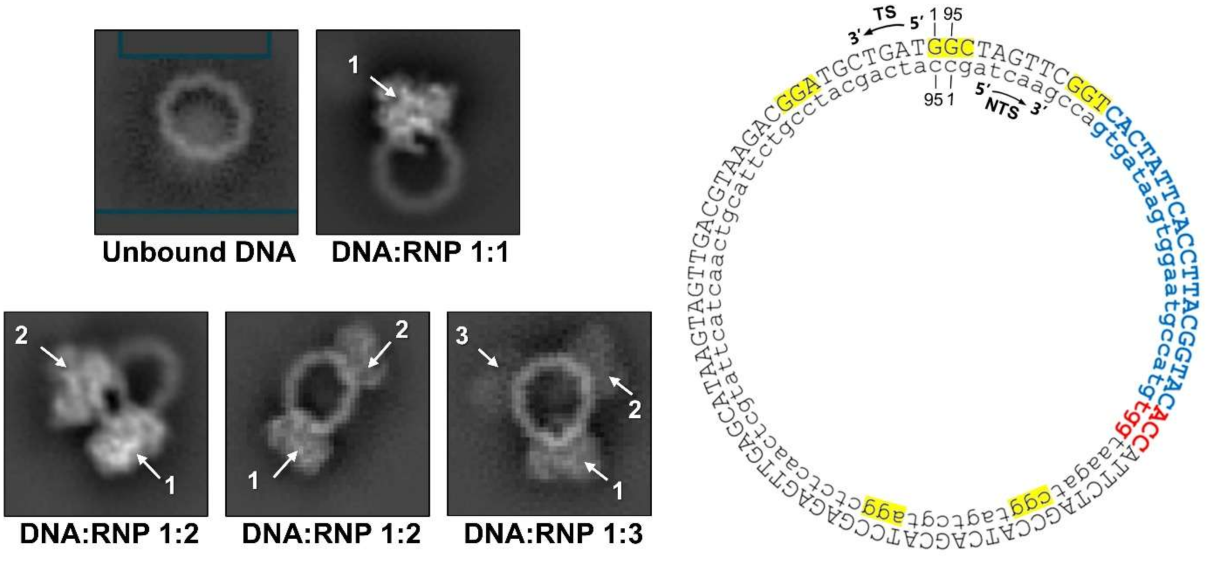


**Figure S6. Heterogeneity of Cas9-dsMC95 complexes revealed by cryo-EM.** Examples of cryo-EM micrograph (left) showing unbound (free) dsMC95 and dsMC95 associated with one to three dCas9 ribonucleoprotein complexes. The unbound (free) dsMC95 likely reflects the weakened binding affinity with Ca9 that results in DNA dissociation between chromatographic purification (Figure S2) and cryo-EM grid preparation. The 1:2 and 1:3 DNA:RNP complexes are most likely off-target binding, as the dsMC95 sequence indivertibly includes multiple "NGG" PAM for SpyCas9 (yellow highlighted on dsMC95 sequence, left) that are not followed by the cognate protospacer (shown in blue). The free dsMC95 and the multiple dCas9-bound dsMC95 "off-target" states would not impact the 1:1 complex analysis presented in this work, as the maps of the 1:1 complex provided sufficient resolution to build the RNA/DNA hybrid in addition to the DNA PAM. However, they are interesting subjects for further investigation.

### Supplementary Figure S7

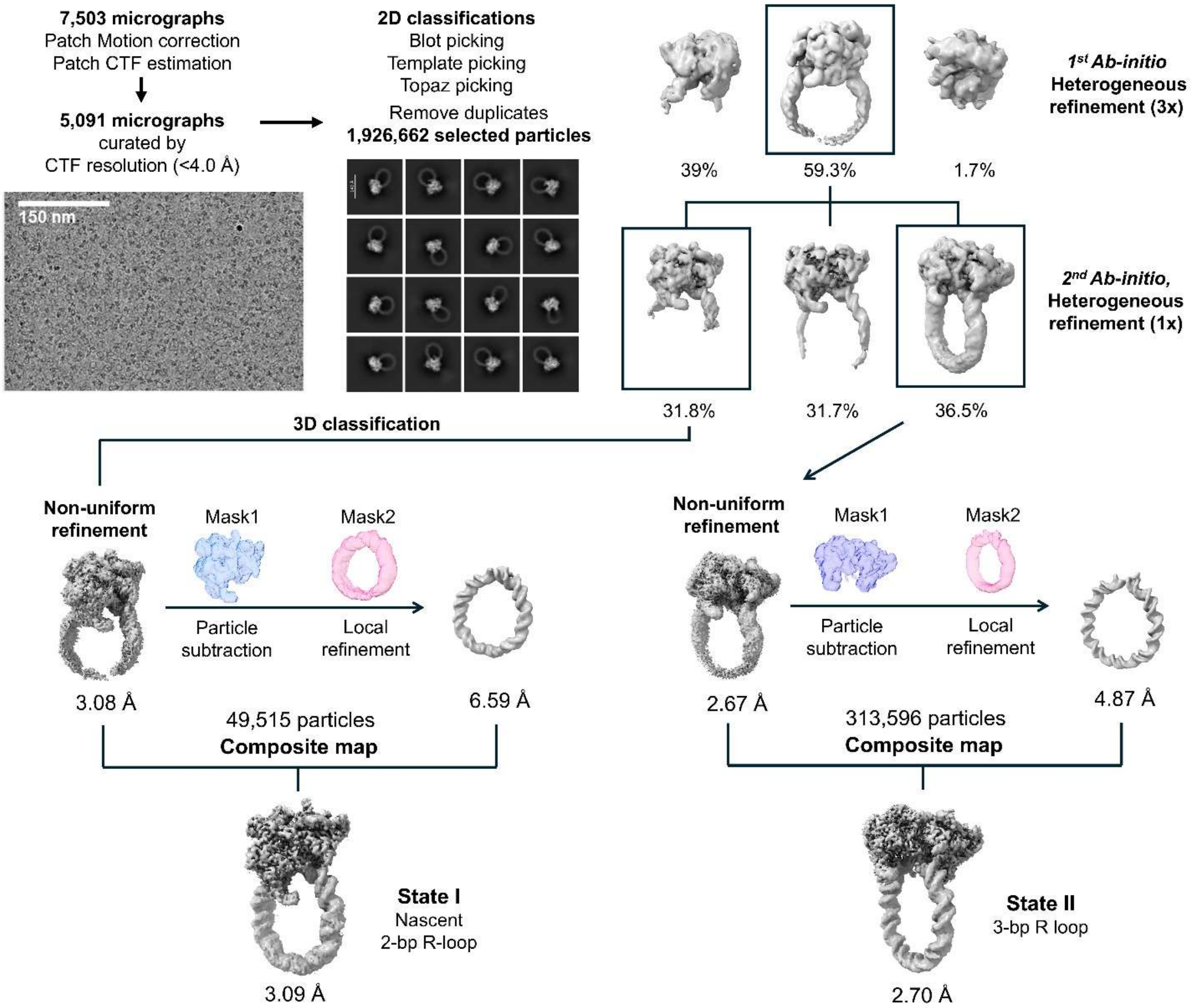


**Figure S7. Cryo-EM data processing workflow.** The pipeline illustrates particle selection from 7,503 micrographs, followed by 2D classification and iterative rounds of *Ab-initio* and heterogeneous refinement. Through non-uniform and local refinement (using particle subtraction), two distinct functional states were resolved as State I and State II described in the main text.

### Supplementary Figure S8

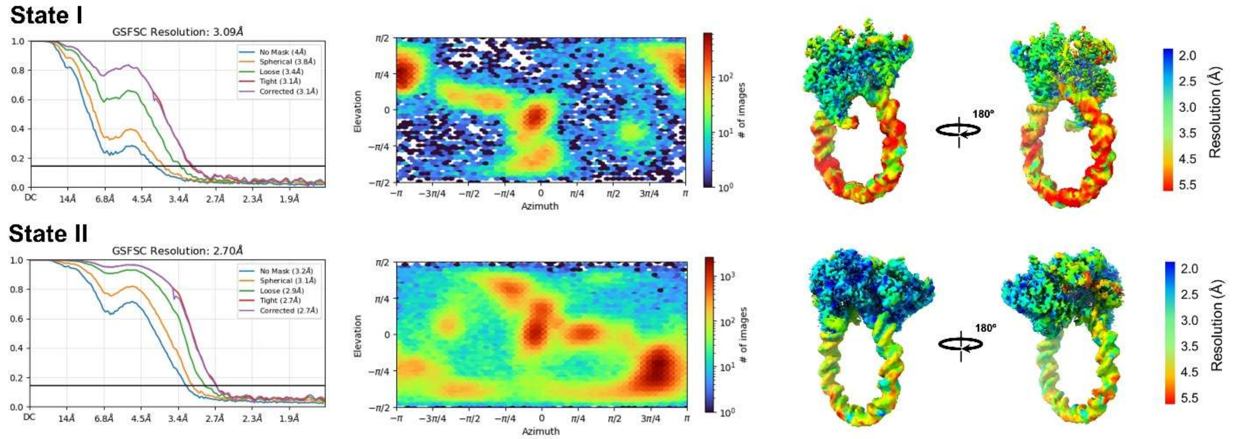


**Figure S8. Validation of the reconstructed cryo-EM maps.** Top and bottom rows correspond to State I and State II, respectively. (Left) Gold-standard FSC curves indicating global resolution at the 0.143 criterion. (Middle) Orientation Distribution Maps showing the orientation coverage of particles. (Right) Cryo-EM density maps colored by local resolution, highlighting the well-resolved core and the more heterogeneous peripheral DNA regions.

**Supplementary Figure S9**

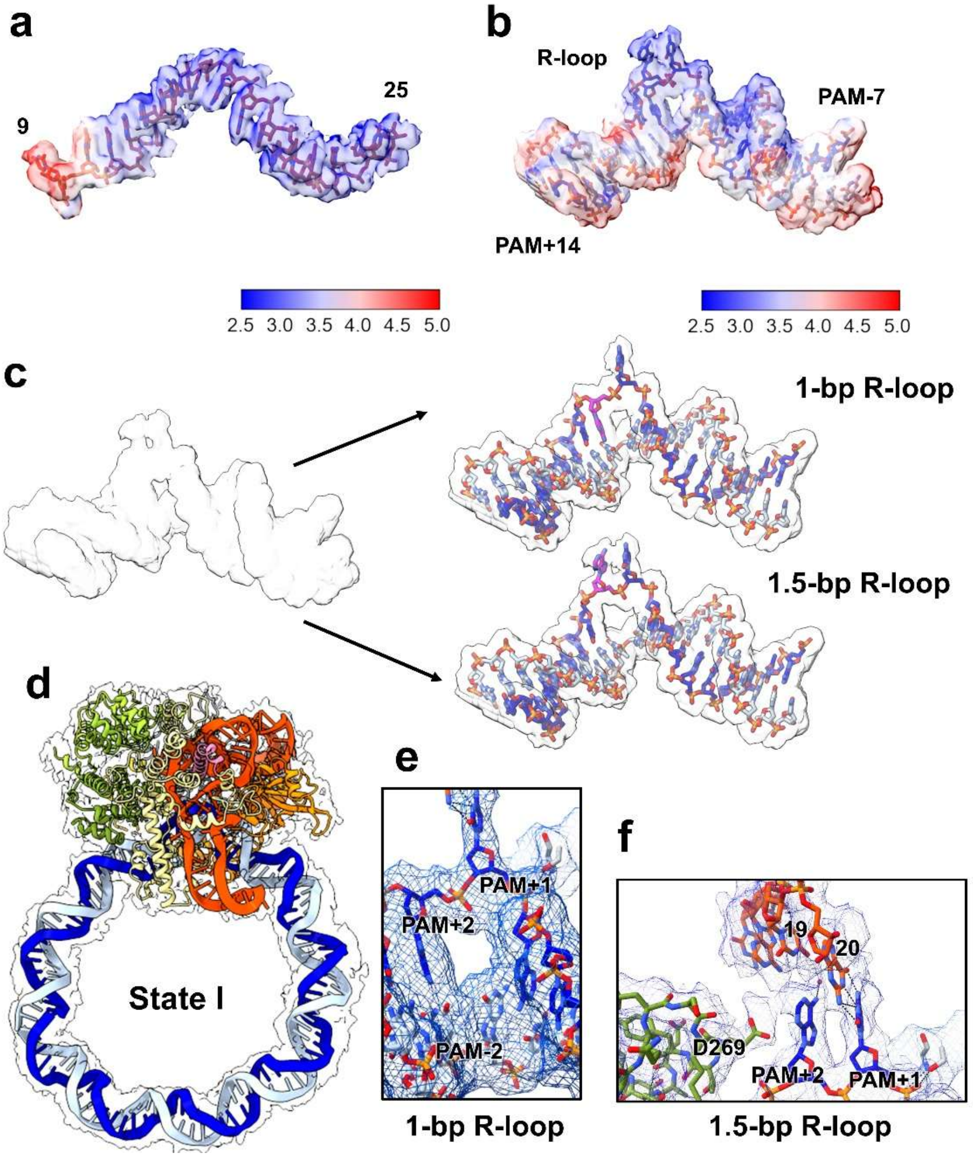


**Figure S9. Cryo-EM model-to-map fitting of State I. a,** Local resolution maps of sgRNA (residues 9–25) in State I. The region shows uniformly high resolution, except for the terminal residue at position 9. **b**, Local resolution maps of dsMC95 spanning from PAM-7 to PAM+14 (see Figure S1b for position designation). The R-loop segment exhibits high resolution, enabling model building of the unwound DNA and subsequently the entire 95-bp DNA. **c**, Coexisting early R-loop intermediates within State I. The map at the R-loop initiation site (left) shows densities consistent with two coexisting sub-states (right). As shown in the map and atomic model overlays, the 1-bp state (top right) shows the PAM+2 nucleotide paired with the target DNA strand, whereas in 1.5-bp (bottom right) shows the PAM+2 nucleotide flipped to engage the sgRNA. **d**, Overlays of cryo-EM density maps and atomic models for State I, with the density shown in white and overlaid with the atomic model (contour level 1.2). **e**, Cryo-EM map and atomic model of the 1-bp R-loop showing good agreement. **f**, Cryo-EM map and atomic model of the 1.5-bp R-loop showing good agreement.

**Supplementary Figure S10**

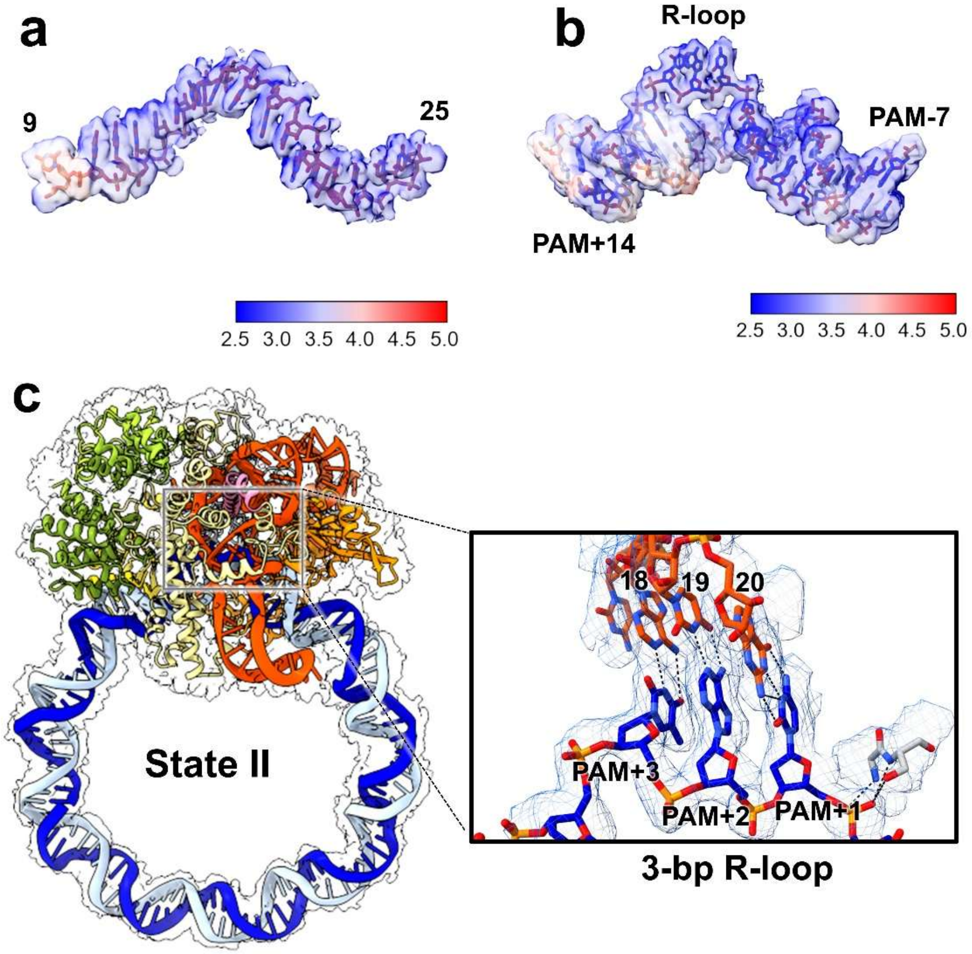


**Figure S10. Cryo-EM model-to-map fitting of state II. a**, Local resolution maps of sgRNA (residues 9–25) in State II. The region shows uniformly high resolution, except for the terminal residue at position 9. **b**, The R-loop region exhibits high resolution, enabling model building of the unwound DNA and subsequently the entire 95-bp DNA. **c**, Overlays of cryo-EM density maps and atomic models for State II, with the density shown in white and overlaid with the atomic model (contour level 2.0). The magnified region (boxed) shows the map (blue mesh) and model overlay spanning the PAM+1 to PAM+3 R-loop.

**Supplementary Figure S11**

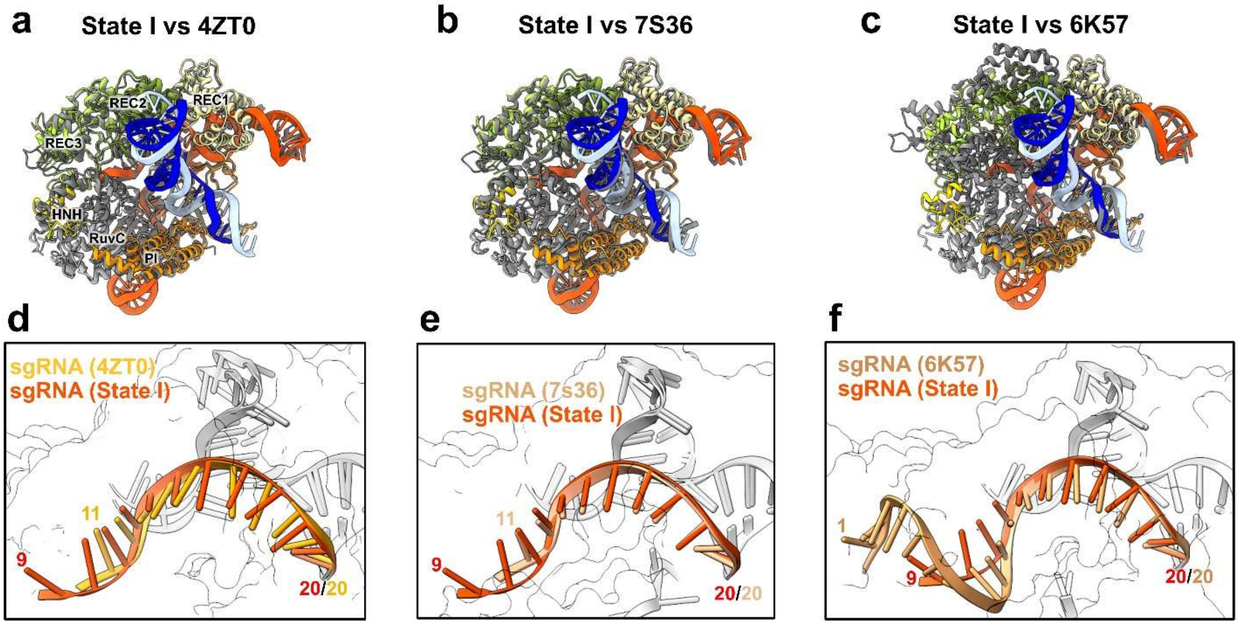


g

| PDB ID | Complex | DNA Type | R-loop | Structural Characteristics | sgRNA @15–20 P RMSD (Å) | Cas9 Cα RMSD (Å) |
|---|---|---|---|---|---|---|
| 13MD, 35SO | Ternary | dsMC95 | 1, 1.5-bp | Flipped PAM+1/+2; poised for R-loop formation | N/A | N/A |
| 4ZT0 | Binary | N/A | N/A | Surveillance-ready state | 0.67 | 1.31 (938 pairs) |
| 7S36 | Ternary | Linear | 0-bp | Flipped PAM+1/+2; non-poised | 0.73 | 1.22 (1,064 pairs) |
| 6K57 | Ternary | Linear | 20-bp | Full R-loop formation | 0.63 | 1.21 (744 pairs) |

**Figure S11. Structural comparison of State I with Cas9 ternary complexes of linear substrates. a–c**, Superposition of State I with Cas9 complexes representing distinct functional states: the surveillance-ready binary complex (PDB: 4ZT0, a), a ternary complex with a 0-bp R-loop (PDB: 7S36, b), and a fully formed 20-bp R-loop state (PDB: 6K57, c). **d–f**, Close-up views of the sgRNA guide showing good alignment from nucleotides 15–20 across the three structures. **g**, Summary of comparisons across the structures presented in **a–c**.

**Supplementary Figure S12**

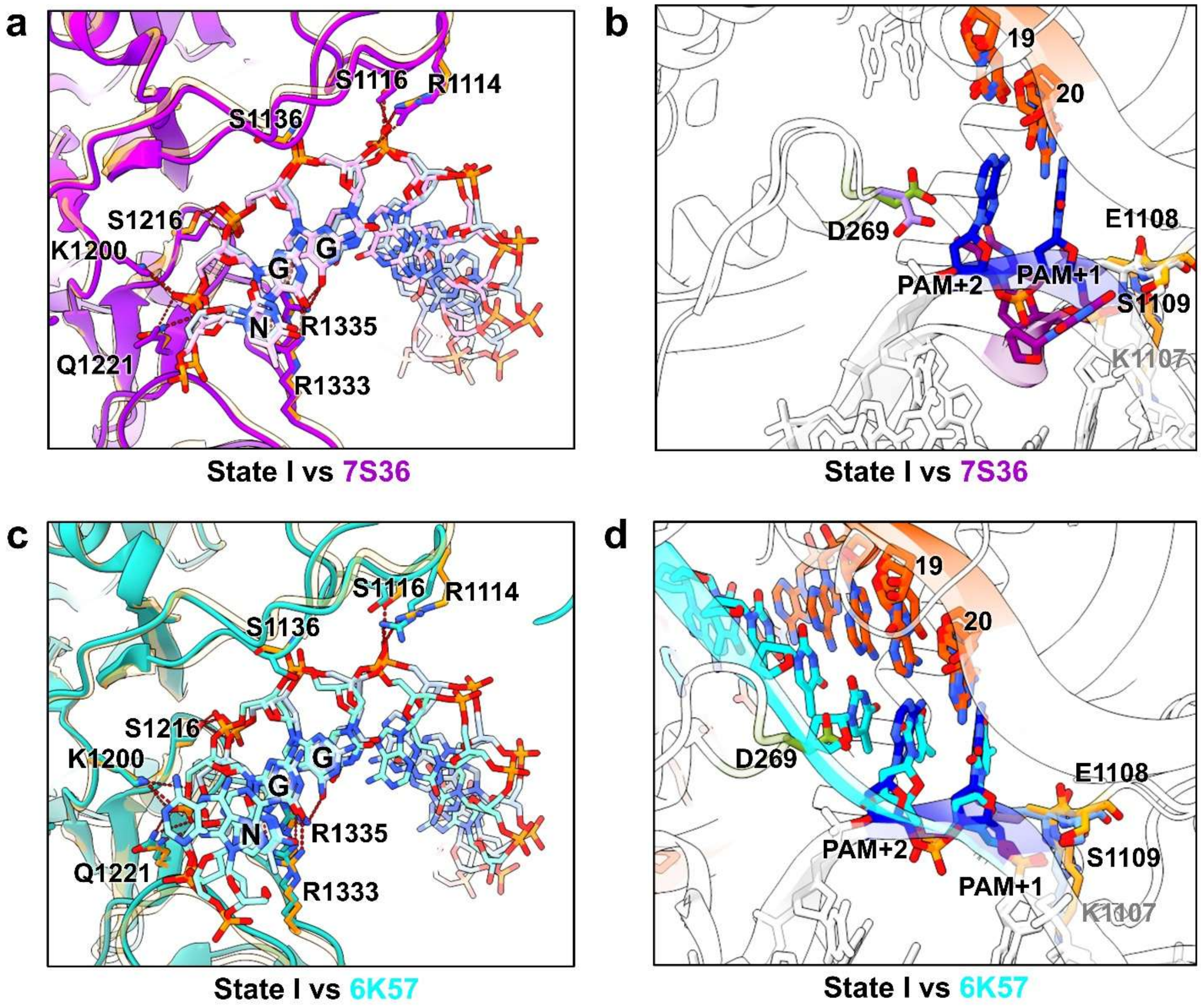


**Figure S12. PAM recognition and phosphate lock in State I. a**, Structural comparison of the PAM-interaction region between State I (wheat) and the "0-bp-linear R-loop" structure of 7S36 (purple). In both structures, side chains of R1333 and R1335 form two hydrogen bonds with the bases in the PAM, indicating that PAM recognition is conserved. **b**, Comparison of the phosphate lock and R-loop initiation region between State I (wheat) and 7S36 (purple). The phosphate lock residue K1107, together with S1109 and E1108, forms a conserved interaction network with the phosphate backbone at the PAM+1 position. **c**, Structural comparison of the PAM-interaction region between State I (wheat) and the "20-bp-linear R-loop" structure of 6K57 (cyan) showing a high degree of alignment. **d**, Comparison of the phosphate lock and R-loop initiation region between State I (wheat) and 6K57 (cyan). The interaction network involving K1107, S1109, and E1108 is maintained, and the R-loop segment (PAM+1 and PAM+2) shows no significant conformational deviation, supporting that State I represents a canonical R-loop initiation geometry.

**Supplementary Figure S13**

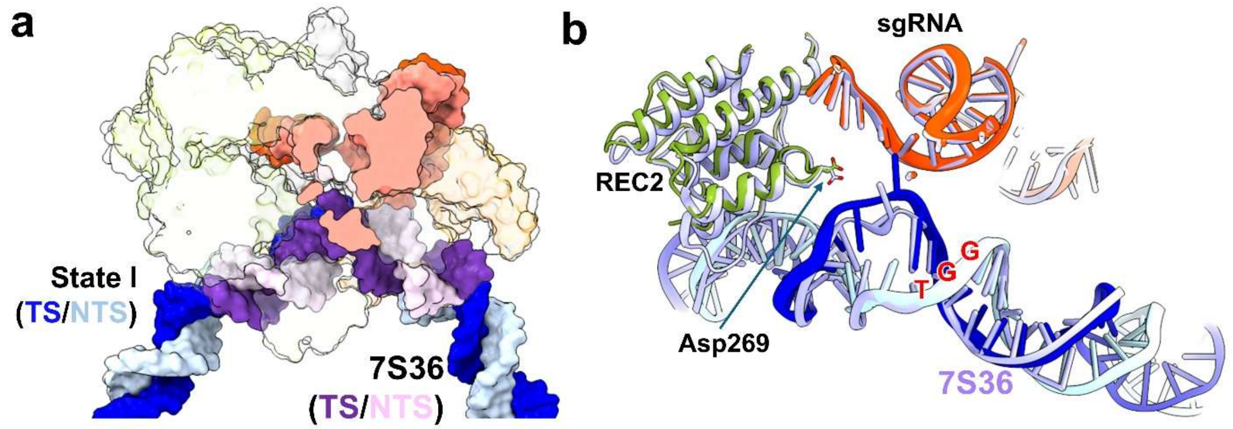


**Figure S13. Structural comparison between State I and the "0-bp-linear R-loop" structure 7S36. a**, Map superimposition of State I (colored by domain) and 7S36 shows very good alignment between the DNA. **b**, Alignment of the atomic models shows very good overlay of the RNA guide (red), the DNA (blue and grey), and the REC2 domain (green and grey) including D269.

### Supplementary Figure S14

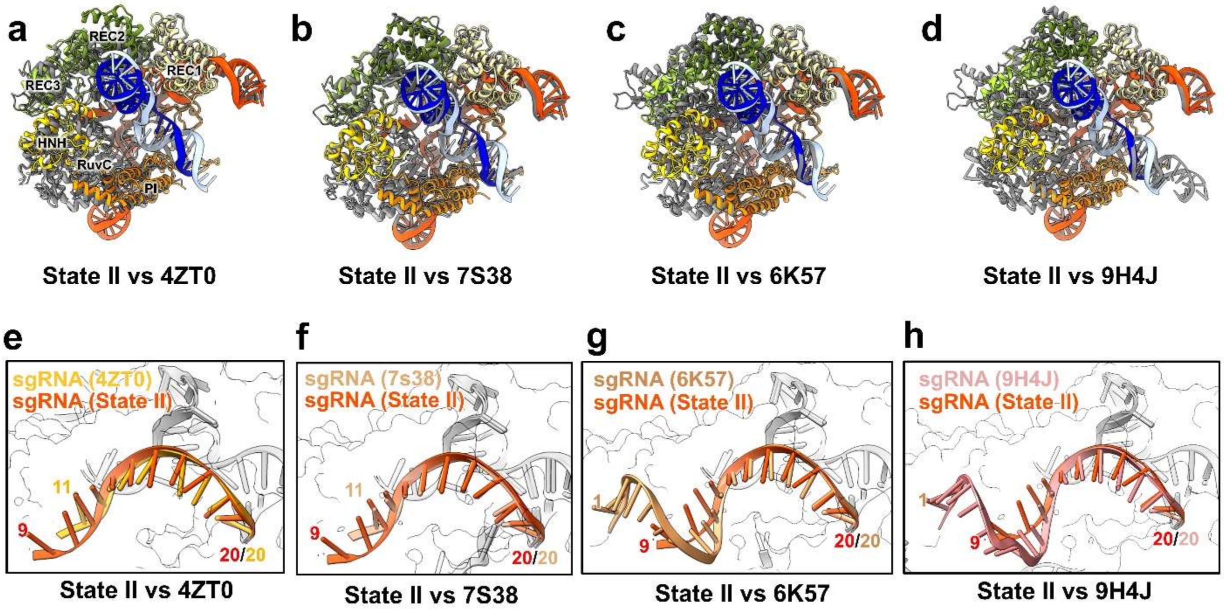


**i**

| PDB ID | Complex | DNA Type | R-loop | Structural Characteristics | sgRNA @15–20 P RMSD (Å) | Cas9 Cα RMSD (Å) |
|---|---|---|---|---|---|---|
| **35SP** | Ternary | dsMC95 | 3-bp | Flipped PAM+1/+2; poised for R-loop formation | N/A | N/A |
| **4ZT0** | Binary | N/A | N/A | Surveillance-ready state | 0.69 | 1.25 (742 pairs) |
| **7S38** | Ternary | Linear | 3-bp | Cross-linking PAM recognition region | 0.59 | 1.23 (1,133 pairs) |
| **6K57** | Ternary | Linear | 20-bp | Full R-loop formation | 0.78 | 1.14 (777 pairs) |
| **9H4J** | Ternary | -sc MC | Like 20-bp | Complex with 126-bp negative sc circular DNA | 1.22 | 0.70 (814 pairs) |

**Figure S14. Structural comparison of State II with other Cas9 ternary complexes. a–d**, Structural superposition of State II with Cas9 complexes representing distinct functional stages: the surveillance-ready binary complex (PDB: 4ZT0, a), a 3-bp R-loop ternary complex assembled with linear DNA (PDB: 7S38, b), a 20-bp R-loop ternary complex assembled with linear DNA (PDB: 6K57, c), and a fully formed R-loop structure assembled on a 126-bp negatively supercoiled circular DNA substrate (PDB: 9H4J, d). **e–h**, Enlarged views of the sgRNA guide region showing very good alignment between positions 15–20. **i**, Summary of comparison across the structures presented in **a–d**.

### Supplementary Figure S15

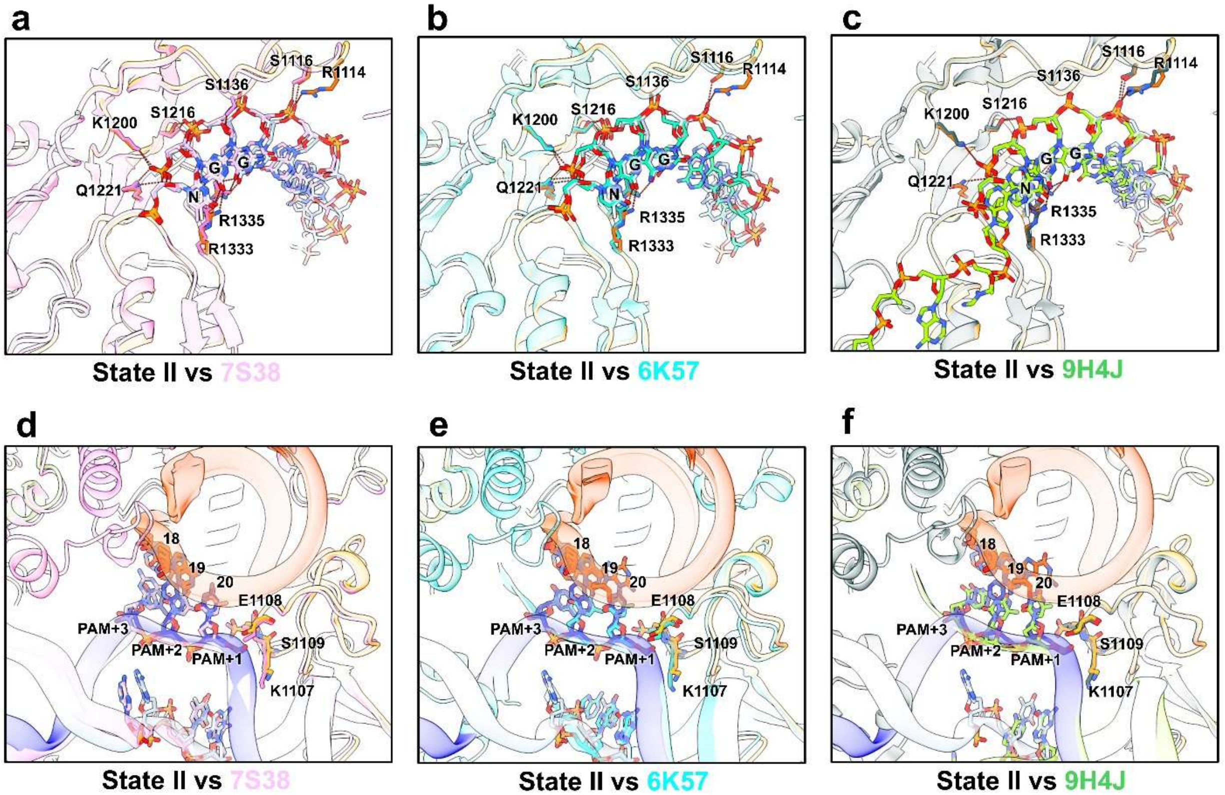


**Figure S15. PAM recognition and phosphate lock in State II. a–c**, Structural comparison of the PAM-interacting region between State II (wheat) and previously reported Cas9 structures 7S38 (pink), 6K57 (cyan), and 9H4J (green). Identical interactions are observed between side chains of R1333 and R1335 and bases in the PAM. **d–f**, Close-up views of the phosphate lock and the R-loop initiation region in comparison between State II and 7S38 (pink), 6K57 (cyan), and 9H4J (green). The phosphate lock residue K1107, together with E1108 and S1109, forms a conserved interaction network with the phosphate backbone at the PAM+1 position. In d–f, State II sgRNA is shown in orange, the target strand (TS) in blue, the non-target strand (NTS) in light blue, and Cas9 in wheat.

**Supplementary Figure S16**

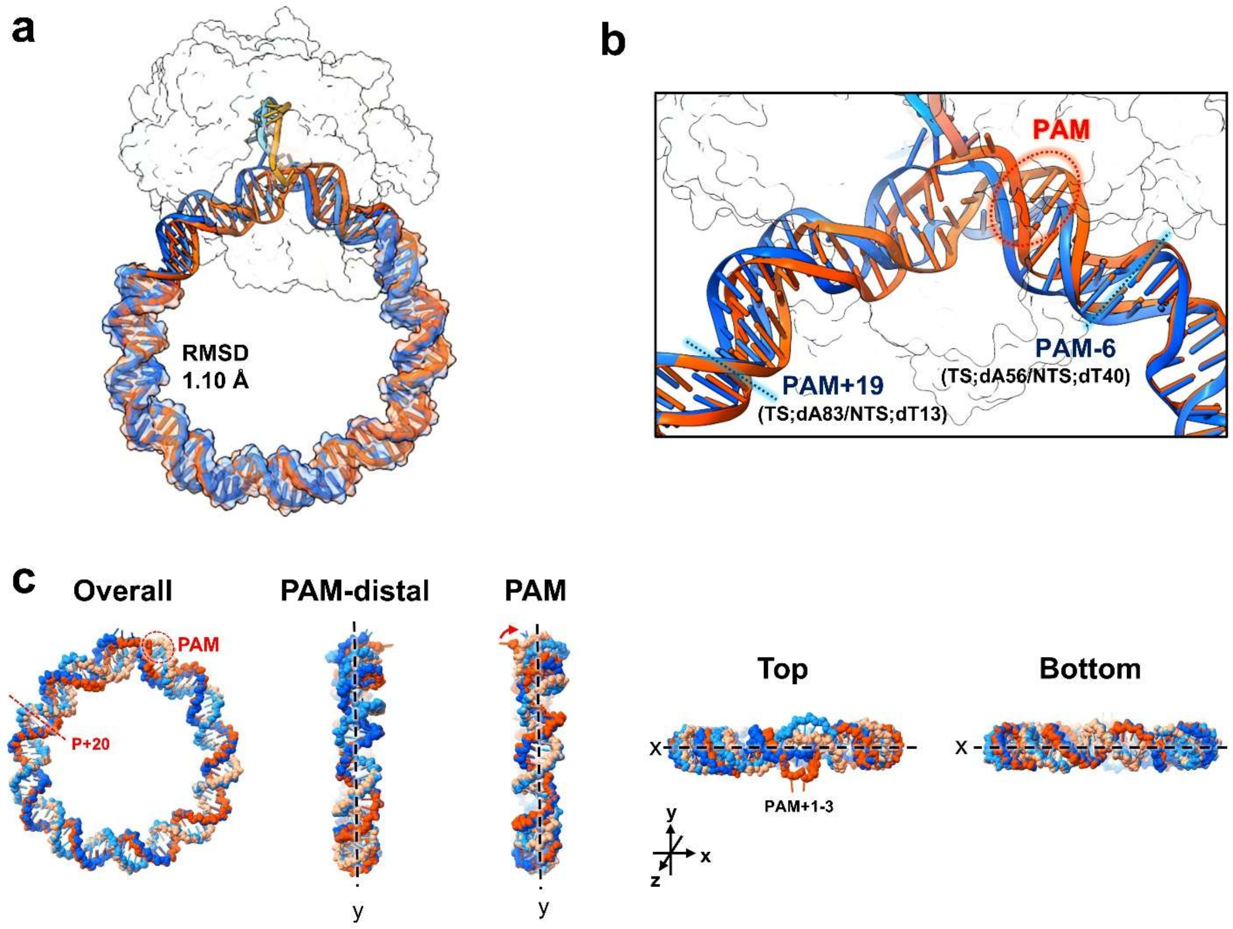


**Figure S16: Superimposition of State I and State II based on the dsMC95 DNA. a**, Overall structural superposition of State I (orange) and State II (blue) aligned on the dsMC95 DNA backbone. The transparent surface indicates the 67-bp region exhibiting an RMSD of 1.1 Å. **b**, Close-up view of the remaining 28-bp region showing DNA deformation. Structural divergence is localized between PAM−6 and PAM+19, where R-loop expansion induces substantial backbone shifts. **c**, Superposition of DNA from State I and State II. Rotation about the y-axis reveals a slight bending transition from State I to State II (red arrow), while rotation about the x-axis shows twisting centered at the PAM+1-3 region as R-loop expands.

**Supplementary Figure S17**

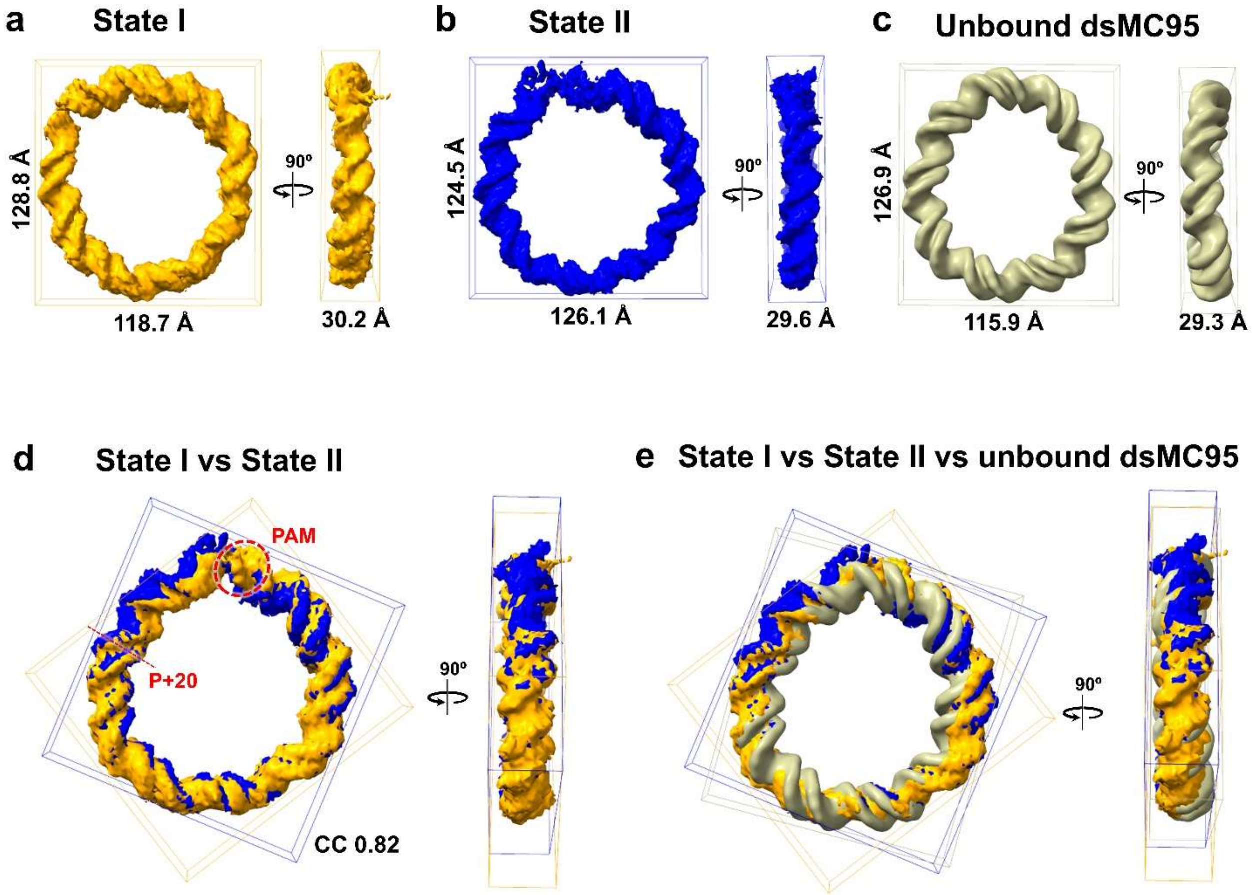


**Figure S17: Comparison of dsMC95 density maps in the Cas9-bound states and the unbound (free) state. a-c**, Cryo-EM density maps of dsMC95 in Cas9-bound State I (orange) and State II (blue) as well as the unbound state (khaki), with boxes showing dimensions estimated using ChimeraX. **d**, Superposition of dsMC95 maps from Cas9-bound State I and State II. **e**, Superposition of dsMC95 maps derived from State I and State II with the unbound dsMC95. All maps are displayed as density surfaces and aligned based on the DNA ring.

**Supplementary Figure S18**

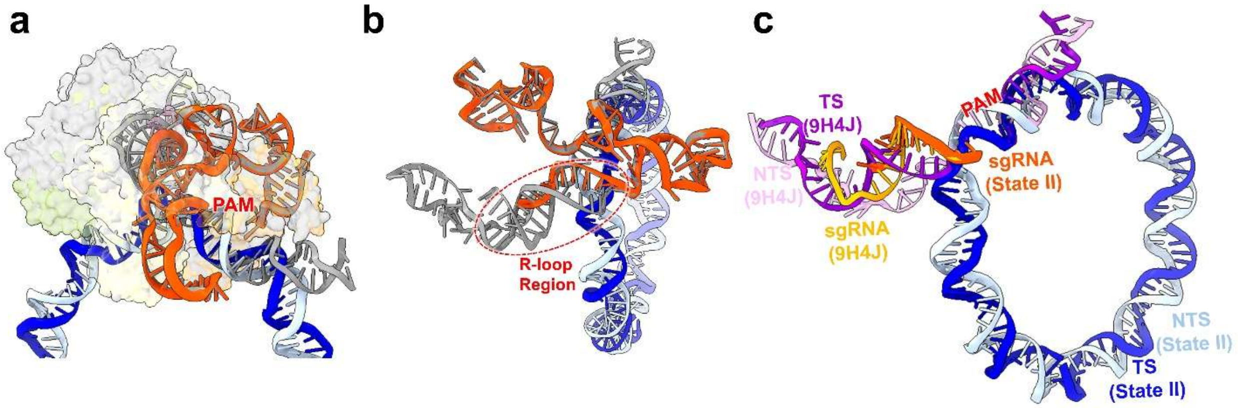


**Figure S18: Structural comparison between Cas9-bound dsMC95 State II and the negative supercoiled DNA complex (PDB 9H4J). a**, Overall structural superposition. The PAM-interacting region aligns well, but differences are observed in the REC2, REC3, and HNH domains. **b**, Comparison of the sgRNA scaffold shows very good alignment. **c**, Structural comparison of DNA. The RNA-guide and the PAM align very well. However, with a full R-loop formed, the overall DNA in 9H4J exhibits opposite bending orientation and twist relative to that of State II, which is stalled at with a 3-bp R-loop.

**Table S1: Sequences of nucleic acids used in this work.**

| **sgRNA** | **Sequences (5′-3′)** |
|---|---|
| **g20**[(a)] | GUGAUAAGUGGAAUGCCAUGGUUUUAGAGCUAGAAAUAGCAAGUUAAAAUAAGGCUAGUCCGUUAUCAA CUUGAAAAAGUGGCACCGAGUCGGUGCUUUU |
| **DNA**[(b)] | **Sequences (5′-3′)** |
| **A1** | CGATCAAGCCAGTGATAAGTGGAATGCCATGTGGTAAGATCGGTAGTC |
| **A2** | GTAGGCTCTCAACTCGTATTCATCAACTGCATTCTGCCTACGACTAC |
| **B1** | GACTACCGATCTTACCACATGGCATTCCACTTATCACTGGCTTGATCG |
| **B2** | GTAGTCGTAGGCAGAATGCAGTTGATGAATACGAGTTGAGAGCCTAC |
| **A1-40-FAM**[(c)] | CGATCAAGCCAGTGATAAGTGGAATGCCATGTGGTAAGA/FAM_dT/*CGGTAGTC |
| **A1-14-2AP**[(d)] | CGATCAAGCCAGTGATA/2AP/**GTGGAATGCCATGTGGTAAGACGGTAGTC |

(a) Single-stranded guide underlined.
(b) Protospacer sequences colored blue.
(c) A fluorescent dT (6-FAM) substitution is installed and is indicated by the bolded "/dT/".
(d) A 2-aminopurine substitution is installed and is indicated by the bolded "/2AP/".

**Supplementary Table S2: Cryo-EM data collection, refinement, and validation statistics for dCas9-bound dsMC95 complexes.**

| | 1-bp & 1.5-bp R-loop dCas9-sgRNA-dsMC95 | 3-bp R-loop dCas9-sgRNA-dsMC95 |
|---|---|---|
| **PDB entry** | 13MD | 35SP |
| **EMDB entry** | | |
| Composite map | EMD-77151 | EMD-77167 |
| Consensus map | EMD-77147 | EMD-77165 |
| Focused map (dsMC95) | EMD-77150 | EMD-77166 |
| **Data collection and processing** | | |
| Magnification | 105,000× | 105,000× |
| Voltage (kV) | 300 | 300 |
| Electron exposure (e–/Å$^2$) | 51.3 | 51.3 |
| Defocus range (μm) | -1.0 to -2.4 | -1.0 to -2.4 |
| Pixel size (Å) | 0.85 | 0.85 |
| Symmetry imposed | C1 | C1 |
| Initial particle images (no.) | 1,926,662 | 1,926,662 |
| Final particle images (no.) | 49,515 | 313,596 |
| Map resolution (Å) | 3.09 | 2.70 |
| FSC threshold | 0.143 | 0.143 |
| Map resolution range (Å) | 2–5.5 | 2–5.5 |
| Map sharpening *B* factor (Å$^2$) | -42 | -59.8 |
| **Refinement** | | |
| Initial model used (PDB code) | 6K57 | 6K57 |
| Model resolution (Å) | 3.3 | 2.9 |
| FSC threshold | 0.5 | 0.5 |
| Model resolution range (Å) | N/A | N/A |
| Model composition | | |
| Non-hydrogen atoms | 15,663 | 16,391 |
| Protein residues | 1,205 | 1,300 |
| Nucleotide | 280 | 280 |
| Ligands | 1 | 1 |
| Water | 95 | 158 |
| *B* factors (Å$^2$) | | |
| Protein | 116.3 | 91.2 |
| Nucleotide | 173.2 | 161.4 |
| Ligand | 47.5 | 23.5 |
| Water | 91.1 | 38.4 |
| R.m.s. deviations | | |
| Bond lengths (Å) | 0.004 | 0.004 |
| Bond angles (°) | 0.563 | 0.553 |
| Validation | | |
| MolProbity score | 1.52 | 1.29 |
| Clashscore | 4.56 | 3.94 |
| Poor rotamers (%) | 0.0 | 0.0 |
| Ramachandran plot | | |
| Favored (%) | 95.86 | 97.43 |
| Allowed (%) | 4.14 | 2.57 |
| Disallowed (%) | 0.0 | 0.0 |
